# Seasonal song variation in male Carolina Wrens (*Thryothorus ludovicianus*)

**DOI:** 10.1101/2021.05.05.442787

**Authors:** Holly R. Keating, Dustin G. Reichard

## Abstract

Birdsong’s primary function is attracting and stimulating mates and repelling rivals during the breeding season. However, many species also sing during the nonbreeding season, which raises questions about the function of nonbreeding song and the proximate mechanisms underlying its production. In this study, we compared spectral and temporal measurements from a large sample of breeding (*N* = 267) and nonbreeding songs (*N* = 283) of Carolina Wrens (*Thryothorus ludovicianus*), a nonmigratory, temperate species that sings year-round. We found that breeding songs were longer than nonbreeding songs and had more syllables within each song. Trill rate, the number of notes per syllable, minimum and maximum frequency and frequency bandwidth did not differ detectably between the two seasons. This study is the first to examine seasonal song differences in Carolina Wrens and provides a basis for future investigations into the drivers behind this seasonal variation.

---

The majority of research on avian vocal behavior has focused on male songs during the breeding season in north temperate climates (Catchpole and Slater 2008). This bias has led to the assumption that most songbirds primarily sing to attract mates and repel rivals during breeding with songs being replaced by calls or silence for the remainder of the year. However, many species of both temperate and tropical songbirds are known to sing outside the breeding season (reviewed in Gahr 2020), including migratory species that sing on their wintering grounds (Sorensen et al. 2016). Despite the abundance of nonbreeding songs, the seasonality of song production and structure has been studied quantitatively in very few species. In those species that have been studied, seasonal changes in song complexity and structure such as song length, syllable repetition rates, syllable consistency, and repertoire composition are inconsistent and species specific (Gahr 2020). Expanding our focus to nonbreeding songs will provide a more complete understanding of the proximate mechanisms underlying song production as well as the full array of functions that songs might serve.

Carolina Wrens (*Thryothorus ludovicianus*) are nonmigratory songbirds found across the eastern United States and Mexico that sing during the breeding and nonbreeding seasons (Fig. 1). Unlike most temperate songbirds, Carolina Wrens form long-term pair bonds and both members of the pair engage in territorial defense year-round (Morton and Shalter 1977, Haggerty et al. 2001). Only males sing, and each individual produces a repertoire of 17-55 (mean = 32) unique song types (Fig. 1; Morton 1987). Song functions in territorial behavior, but song production seems to increase at the onset of the breeding season, suggesting an additional function in courtship and mate stimulation (Haggerty and Morton 2020). A function in mate attraction is less clear given that pair bonds can form in the first few months of life before males fully develop a crystallized song (Morton 1982). Despite this persistent vocal behavior, seasonal variation in the structure of Carolina Wren song has yet to be quantified.

**Figure 1.**
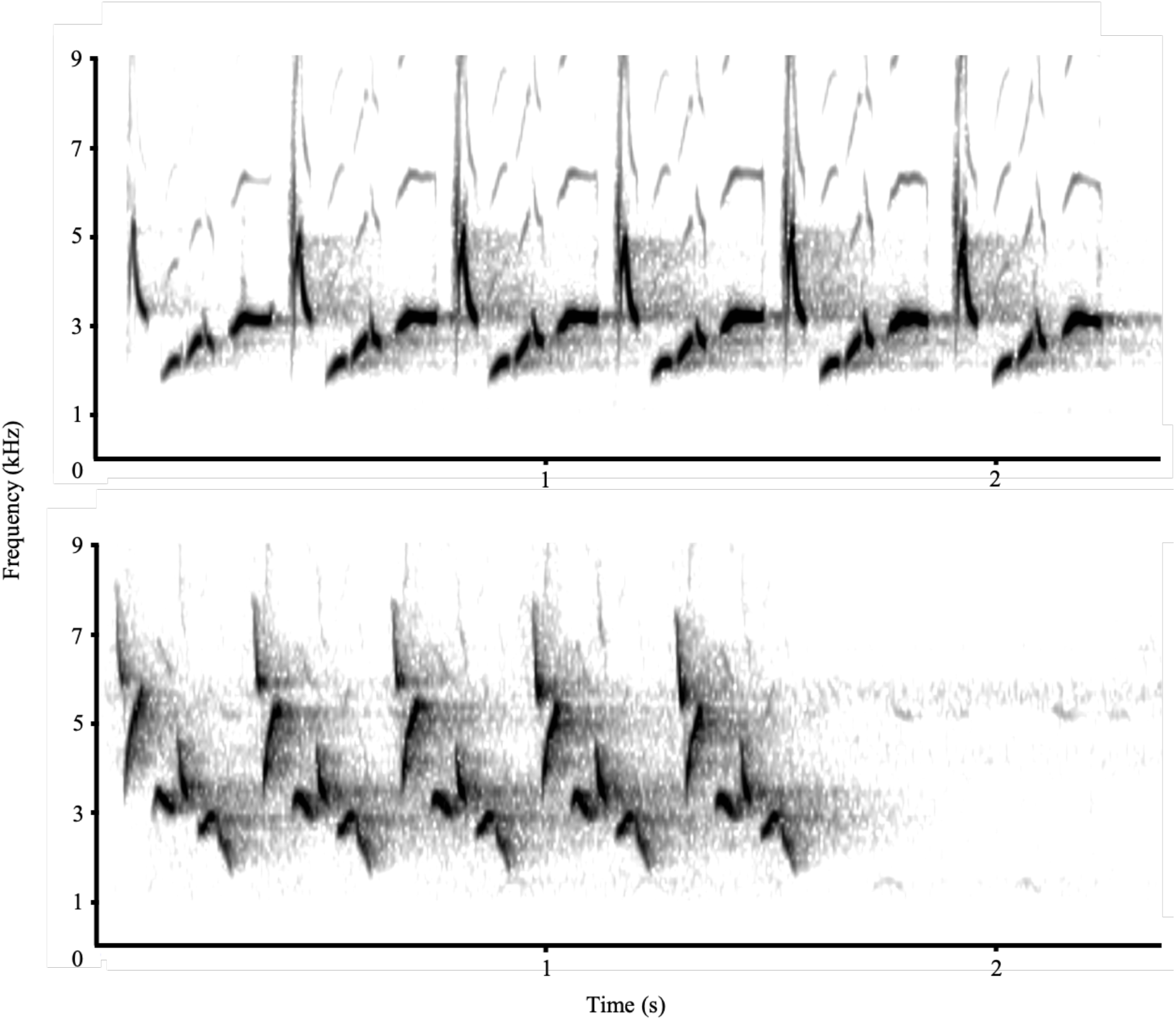
Representative spectrograms of Carolina Wren songs from the breeding (top) and nonbreeding (bottom) seasons. The darkness of the spectrograms does not represent the relative amplitudes of the songs.

In this study, we compared breeding and nonbreeding Carolina Wren songs across a large geographic scale using recordings from online databases. We hypothesized that songs would differ between the breeding and nonbreeding seasons. Specifically, we predicted that breeding season songs would be longer and more complex (more notes/syllable, higher entropy) due to song’s presumed function in courtship and mate stimulation during breeding. In addition, nonbreeding songs may be shorter and less complex due to seasonal reductions in the size of brain nuclei responsible for song production even in species that sing all year (Smith et al. 1995, 1997; but see Leitner et al. 2001, Smulders 2005).

## Methods

### Song Recordings

We obtained recordings of Carolina Wren *(Thryothorus ludovicianus)* songs from the Florida Museum of Natural History (N = 49; WAV format), xeno-canto (N = 124; MP3 format; https://www.xeno-canto.org/), and the Macaulay Library (N = 377; WAV format; https://www.macaulaylibrary.org; see Supplementary Table S1). The recordings were collected between 1954-2020 throughout the Carolina Wren’s range in Canada, the United States, and Mexico (Supplementary Table S1). We inspected each recording for accuracy and quality (signal-to-noise ratio). After removing misidentified and low-quality recordings, we cross referenced the metadata for each recording to remove duplicate songs that may have been uploaded to multiple databases. We also scrutinized the recording dates and locations to prevent the same song type from the same individual from being sampled more than once. A total of 550 unique songs were included in our analysis.

Based on the breeding phenology of Carolina Wrens (Haggerty and Morton 2020), the breeding season encompasses March to August (*N* = 267) and the nonbreeding season is September to February (*N* = 283). In the southern United States, egg laying begins in mid-March and ceases by mid-August, but pairs can continue to feed young as late as early October (Haggerty and Morton 2020). To ensure that our results were not skewed by samples from the transitional months between breeding and nonbreeding, we ran an identical analysis that excluded all recordings collected in February and September (*N* = 106 songs excluded). The results of this analysis were qualitatively identical to our complete analysis included below (Table 1, Supplementary Table S2).

**Table 1.** Results of linear mixed model analysis for song measurements comparing breeding and nonbreeding populations. All means are presented ± 1 SEM. Bolded values identify P < 0.05.

| Song Measurement | Breeding Mean | Nonbreeding Mean | $\chi^2_{df}$ | $P$-value |
| --- | --- | --- | --- | --- |
| **Song Length (sec)** | **1.69±0.02** | **1.56±0.02** | **18.77** | **< 0.001** |
| Maximum Frequency (Hz) | 5483.0±61.24 | 5453.8±59.51 | 0.07 | 0.79 |
| Minimum Frequency (Hz) | 1788.9±16.68 | 1828.1±17.71 | 2.06 | 0.15 |
| Frequency Bandwidth (Hz) | 3694.03±62.22 | 3625.70±60.38 | 0.31 | 0.58 |
| Number of Inflection Points | 94.96±1.72 | 97.95±1.91 | 0.01 | 0.92 |
| Maximum Entropy | 4.90±0.02 | 4.85±0.03 | 1.43 | 0.23 |
| **Number of Syllables/Song** | **4.78±0.09** | **4.26±0.08** | **11.9** | **< 0.01** |
| Number of Notes/Syllable | 4.86±0.07 | 4.92±0.07 | 0.006 | 0.94 |
| Trill Rate (syllables/sec) | 2.79±0.06 | 2.76±0.05 | 0.06 | 0.8 |

### Song Measurements

We used Raven Pro 1.6 (Center for Conservation Bioacoustics, 2011) to measure song length, minimum and maximum frequency, frequency bandwidth, the number of inflection points, maximum entropy, and trill rate. A single observer (HRK) analyzed all songs and was blind to breeding season. We generated spectrograms of each recording (Hann Window, 512 DFT, 93.8 Hz frequency resolution), and used the band limited energy detector to locate high quality songs. Then we randomly chose one song from each recording for analysis.

To minimize human bias, the start and end time for each song was determined by the energy detector, and we used the peak frequency contour function to measure the minimum and maximum frequency and the number of inflection points. The peak frequency contour function traces the peak frequency of a vocalization by measuring individual spectrogram slices through time, which allows for automatic, less biased spectral measurements. The number of inflection points is calculated as the number of times the slope of the peak frequency contour changes signs across the entire song. Maximum entropy was calculated automatically.

We manually counted the number of syllables and notes per syllable for each song and trill rate was calculated as the number of syllables divided by song length. In a few cases (*N* = 30), the energy detector was unable to recognize songs in the recording despite a strong signal-to-noise ratio. When this occurred, we manually determined the start and end time by visually drawing a selection box around the song before collecting the remaining measurements with the peak frequency contour function.

### Statistical Analysis

We conducted linear mixed models using the ‘lme4’ package in R v 4.0.2 (Bates et al. 2015; R Core Team, 2020) to compare breeding and nonbreeding songs. We used model selection by Akaike’s Information Criterion to assess the relative importance of latitude and sampling year as explanatory variables. For each song measurement, a series of models were created that contained the song measurement as the response variable, breeding status as a random factor, and latitude, sampling year, or both as independent factors. The models that included latitude alone were the only models to outperform the null model. We used a likelihood ratio test (LRT) to assess significance by comparing the likelihood of the full model that included latitude to that of a reduced model without breeding season.

## Results

Breeding male Carolina Wrens sang longer songs than non-breeding wrens (Table 1, Fig. 2, *X*^2^_1_ = 18.65, *P* < 0.001) and had more syllables within each song (Table 1, Fig. 2, *X*^2^_1_ = 11.90, *P* < 0.001). There was no difference between breeding and nonbreeding songs in minimum or maximum frequency, frequency bandwidth, the number of notes per syllable, trill rate, the number of inflection points within a song, or maximum entropy (Table 1).

**Figure 2.**
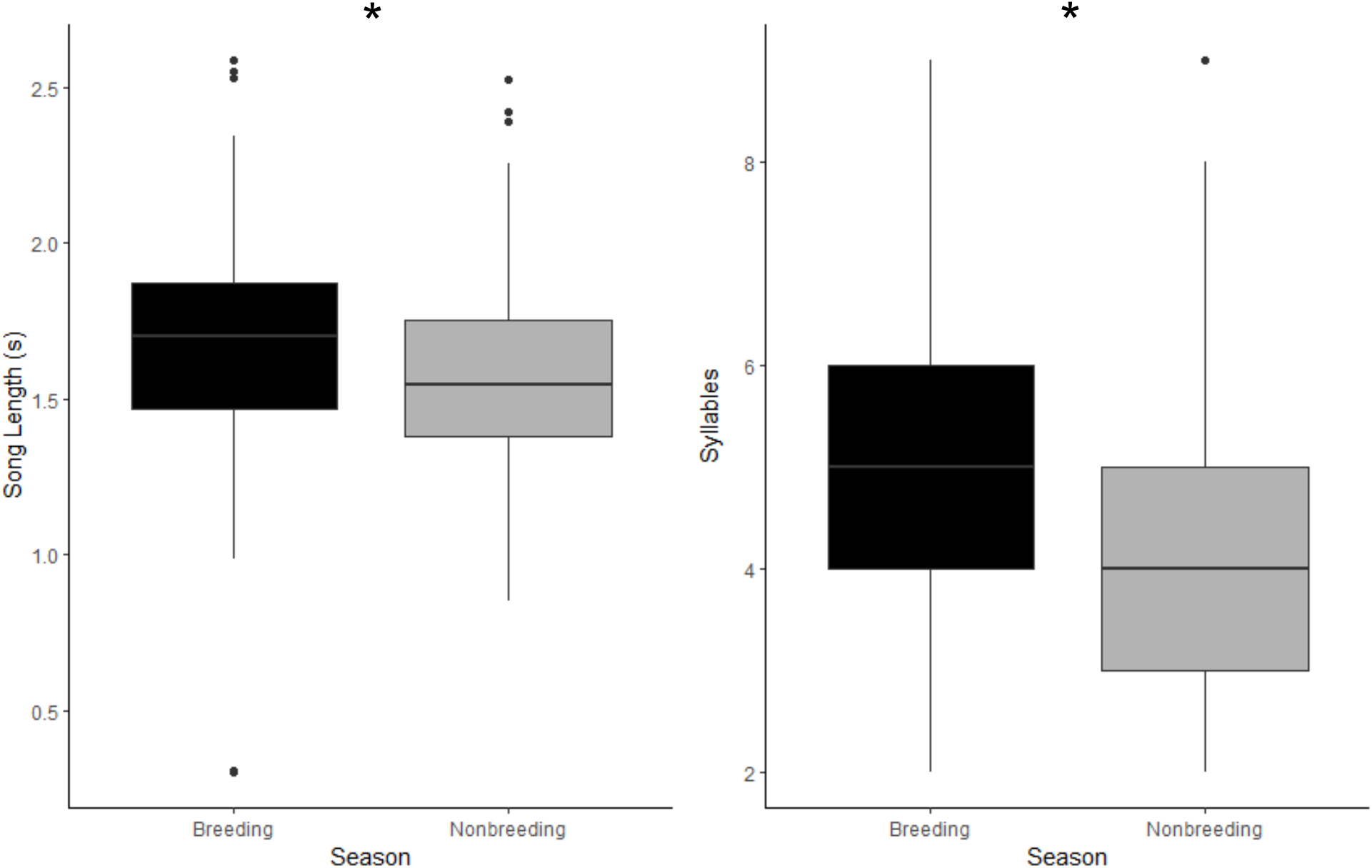
Comparison of song measurements from Carolina Wrens that showed a significant difference between breeding and nonbreeding seasons. Each box represents the interquartile range and median, whiskers represent range of data within 1.5 times the interquartile range, and dots represent data points exceeding that range. \**P* < 0.01

## Discussion

Male Carolina Wrens sang longer songs with more syllables during the breeding season. This result was consistent with our predictions as well as previous seasonal comparisons from other species including both residents (e.g. Nuttall’s White-crowned Sparrows [*Zonotrichia leucophrys nutalli*], Brenowitz et al. 1998; Wild Canaries [*Serinus canaria*], Leitner et al. 2001; Black-Capped Chickadees [*Poecile atricapillus*], Smulders 2005; Rufous-capped Warblers [*Basileuterus rufifrons*], Demko and Mennill 2019) and migrants (Gambel’s White-crowned Sparrows [*Zonotrichia leucophrys gambelii*], Smith et al. 1995). However, we found no detectable seasonal differences in trill rate, the number of notes per syllable, number of inflection points, maximum entropy, or any measures of frequency. This outcome indicates that Carolina Wrens are singing similarly structured songs throughout the year rather than producing more complex songs during the breeding season as we predicted. One limitation of this study was that we were unable to compare the song repertoires of individual males between the breeding and nonbreeding seasons, so it remains unknown whether repertoire composition or the stereotypy of individual song types vary seasonally, which has been observed in other species (Smith et al. 1997, Leitner et al. 2001).

The relative lack of seasonal variation in Carolina Wren song structure may be partially explained by the primary function of song in this species. As a temperate resident, Carolina Wrens defend territories year-round, and a persistent function of song appears to be in territorial behavior (Haggerty and Morton 2020). Maintaining a viable territory is not only important for breeding, but nonbreeding territories are critical for overwinter survival in this species, particularly during harsh winters when food resources become scarce and adult mortality can reach 90% (Morton 1982). This intense nonbreeding territoriality is further supported by the fact that Carolina Wrens ignore dear-enemy effects and engage in more territorial disputes during the nonbreeding season, and males with higher quality territories during harsh winters sing at higher rates than males with low quality territories (Morton and Shalter 1977, Morton 1982, Hyman 2005). Additionally, unpaired females, which do not sing, are unable to maintain territories alone, but solo males can (Morton and Shalter 1977). Collectively, these observations suggest that song’s year-round role in territoriality may be the most important selective force maintaining its consistent structure across the breeding and nonbreeding seasons.

Unlike many temperate songbirds, Carolina Wrens form long term pair bonds that are often established early in life before males develop a crystalized song (Morton and Shalter 1977). Under these conditions, females likely choose mates based on a male’s non-vocal attributes or the quality of his territory, which minimizes the traditional function of song in mate attraction. The presence of long-term genetic monogamy in Carolina Wrens (Haggerty et al. 2001) also makes it unlikely that song functions in attracting extra-pair mates. However, male wrens increase their song rates at the beginning of breeding (Haggerty and Morton 2020), and our study supports a concurrent increase in song length and the number of repetitions of each syllable during the breeding season. Increasing song length and redundancy can increase the detectability of the signal for all potential receivers, which would support functions in both inter and intrasexual communication during breeding (Price 2013). Independent of mate attraction, longer breeding songs may also function in within-pair communication as a contact signal (Baptista 1978) and in stimulating females into reproductive condition (Maney et al. 2007).

The underlying mechanism for the seasonal changes in song length could be due to a variety of factors, including changes in brain nuclei size or shifting hormonal controls. Multiple species of sparrows have divergent song structures in the fall when song control nuclei are significantly smaller (Smith et al 1995, 1997; Brenowitz at al. 1998), but other species such as Black-Capped Chickadees sing longer songs during the breeding season with no seasonal variation in the size of the song system nuclei (Smulders 2005). Male Carolina Wrens have larger song control nuclei than females (Nealen and Perkel 2000), but whether the size of these nuclei vary seasonally is unknown and represents an important topic for future research. Alternatively, changes in sex steroid concentrations or receptor densities and locations could cause seasonal song differences and may or may not work in conjunction with changes in brain nuclei size. Wild canaries, for example, produce longer songs with faster trill rates during the breeding season when plasma testosterone levels are significantly higher, but the size of at least two song system nuclei remains similar between breeding and nonbreeding (Leitner et al. 2001, Voigt and Leitner 2008). Elevated androgen levels appear to consistently increase song rates in seasonally breeding species, but the exact role of androgens in determining song structure, particularly during the nonbreeding season when plasma levels are often low, remains poorly understood (reviewed in Gahr 2020).

The majority of our assumptions about the ultimate and proximate mechanisms underlying birdsong structure and function are derived from studies of breeding songs in temperature species. Nonbreeding songs are common, and their production is often regulated by mechanisms that cannot be easily explained by circulating levels of sex steroids (Gahr 2020). Songs produced outside the breeding season are also more likely shaped by functions in territoriality and competition rather than mate attraction and choice. Expanding our focus to species such as Carolina Wrens, which sing throughout the year, may provide new insights into the mechanisms of song production and the complexity of selective forces that shape song structure.

## Acknowledgments

We thank the Macaulay Library at the Cornell Lab of Ornithology, xeno-canto, and the Florida Museum of Natural History for use of their song recordings. We thank the Ohio Wesleyan University Summer Science Research Program for funding.

---

**Supplementary Table S1.**
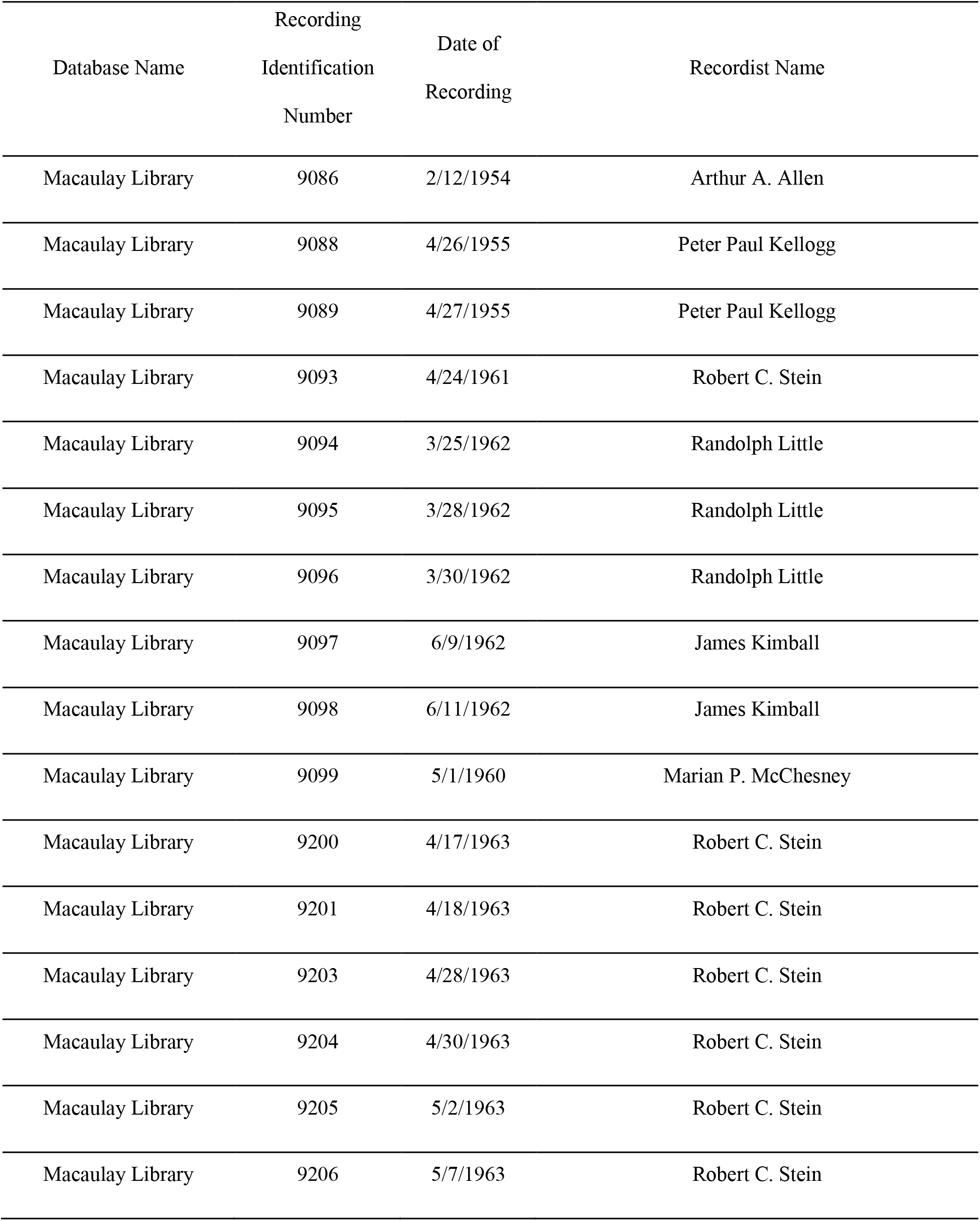
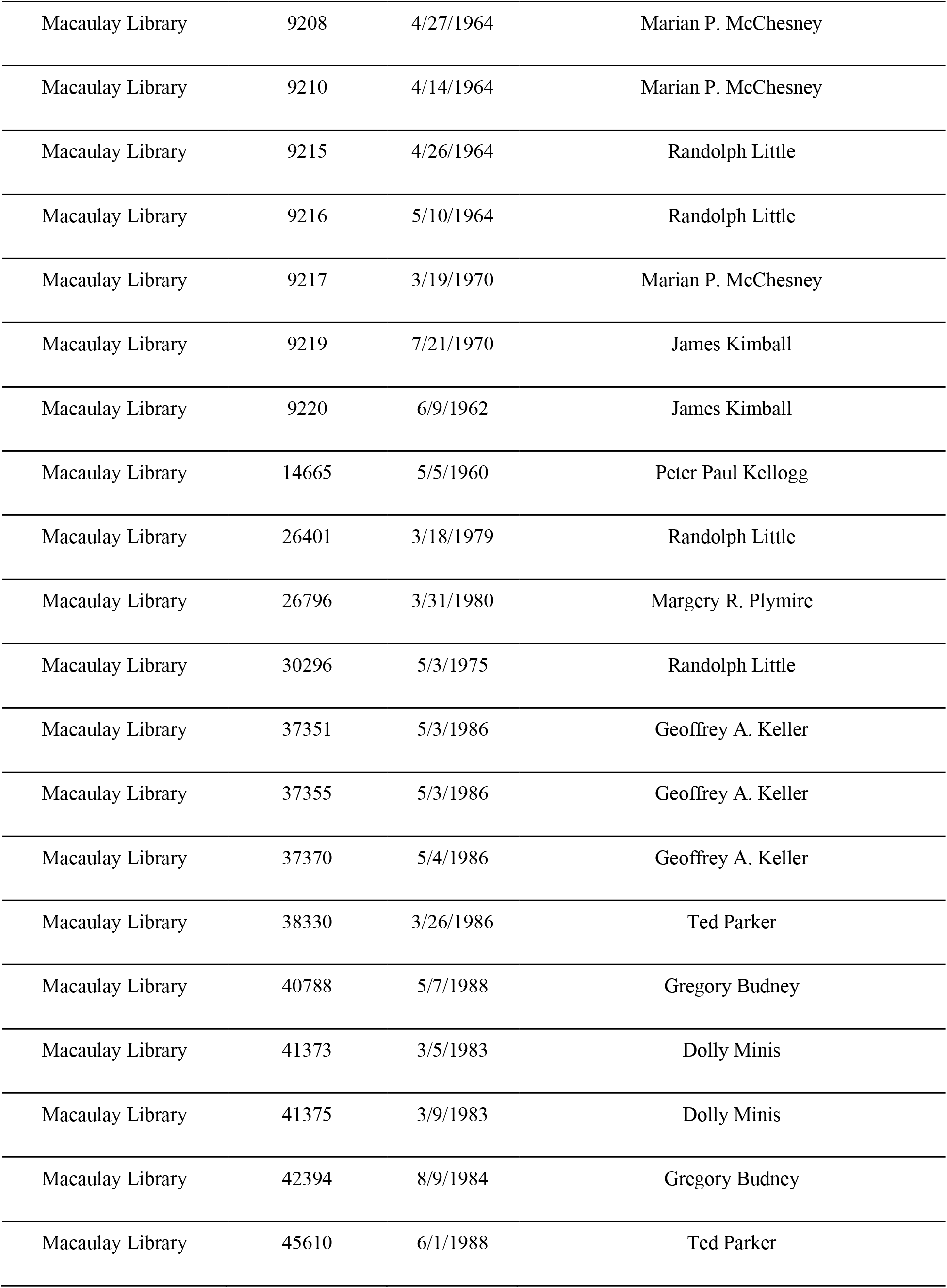
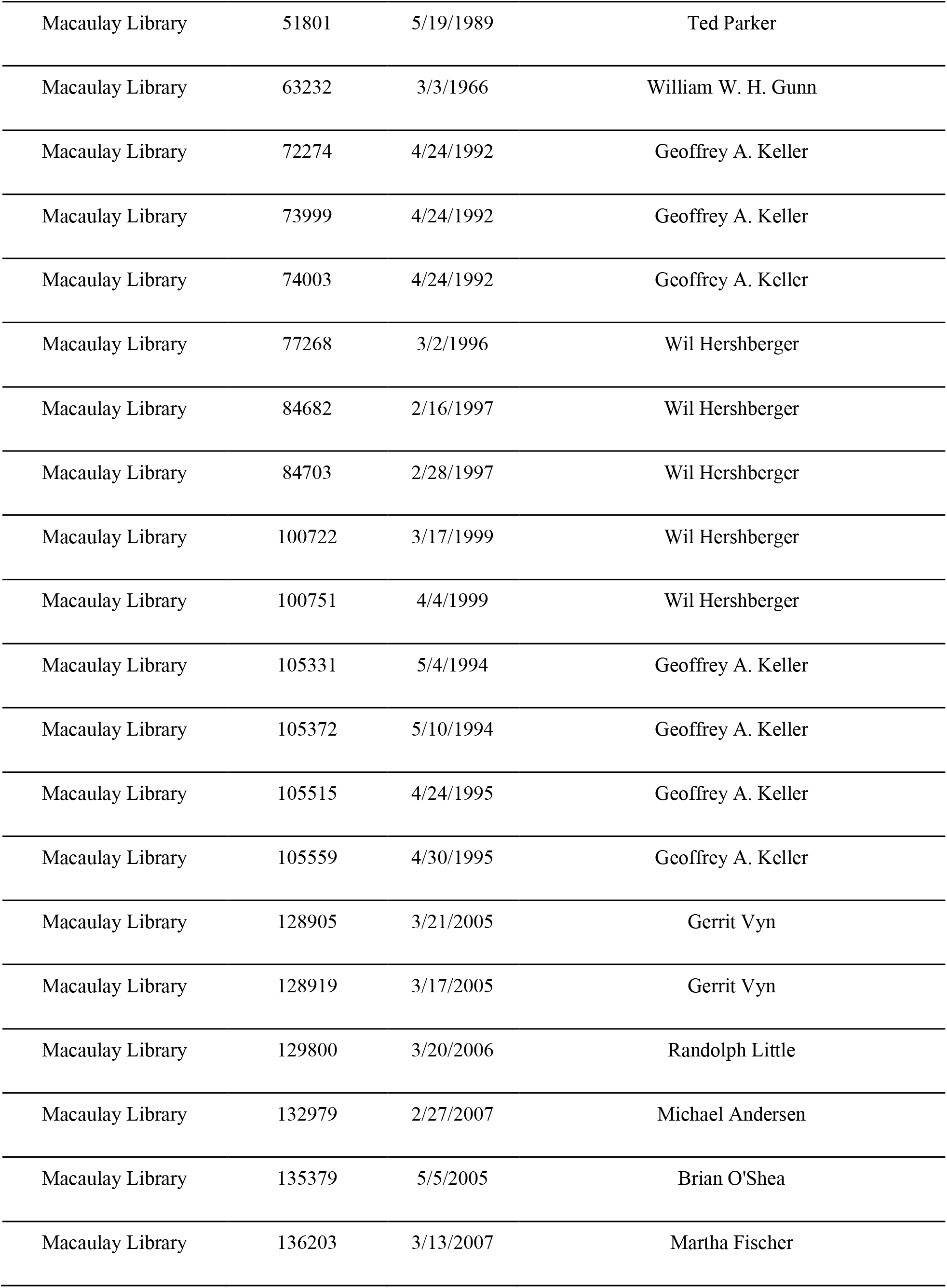
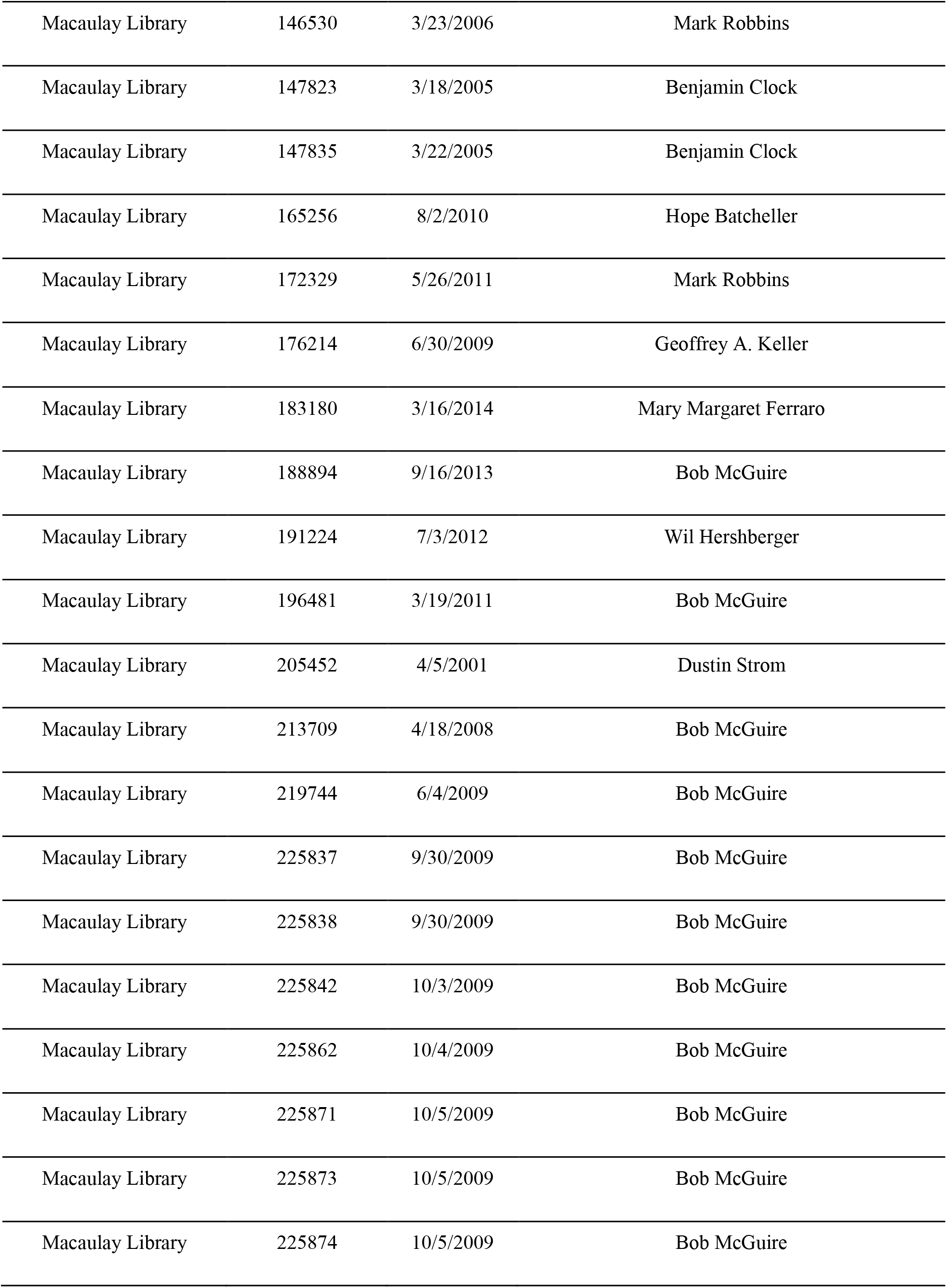
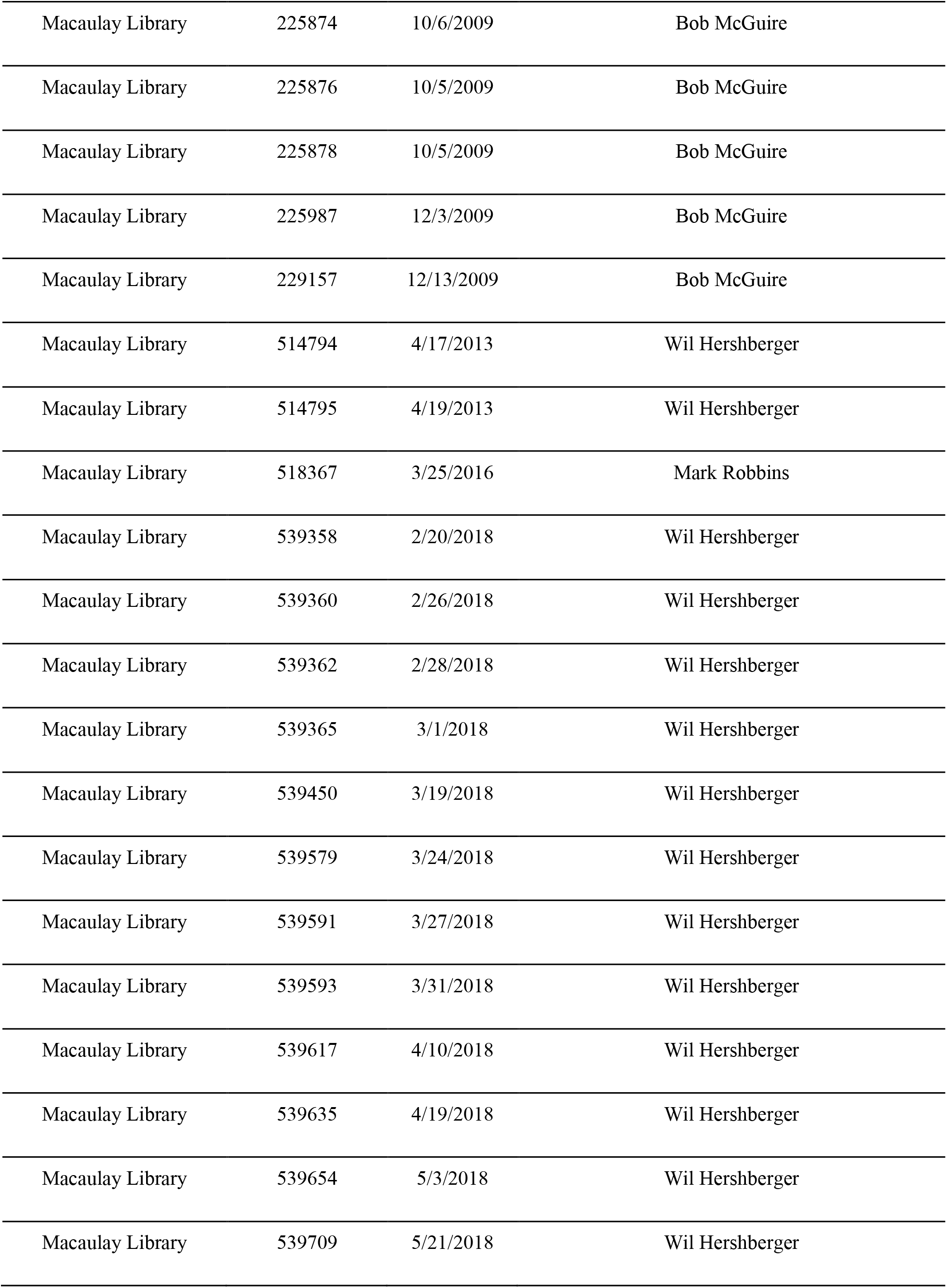
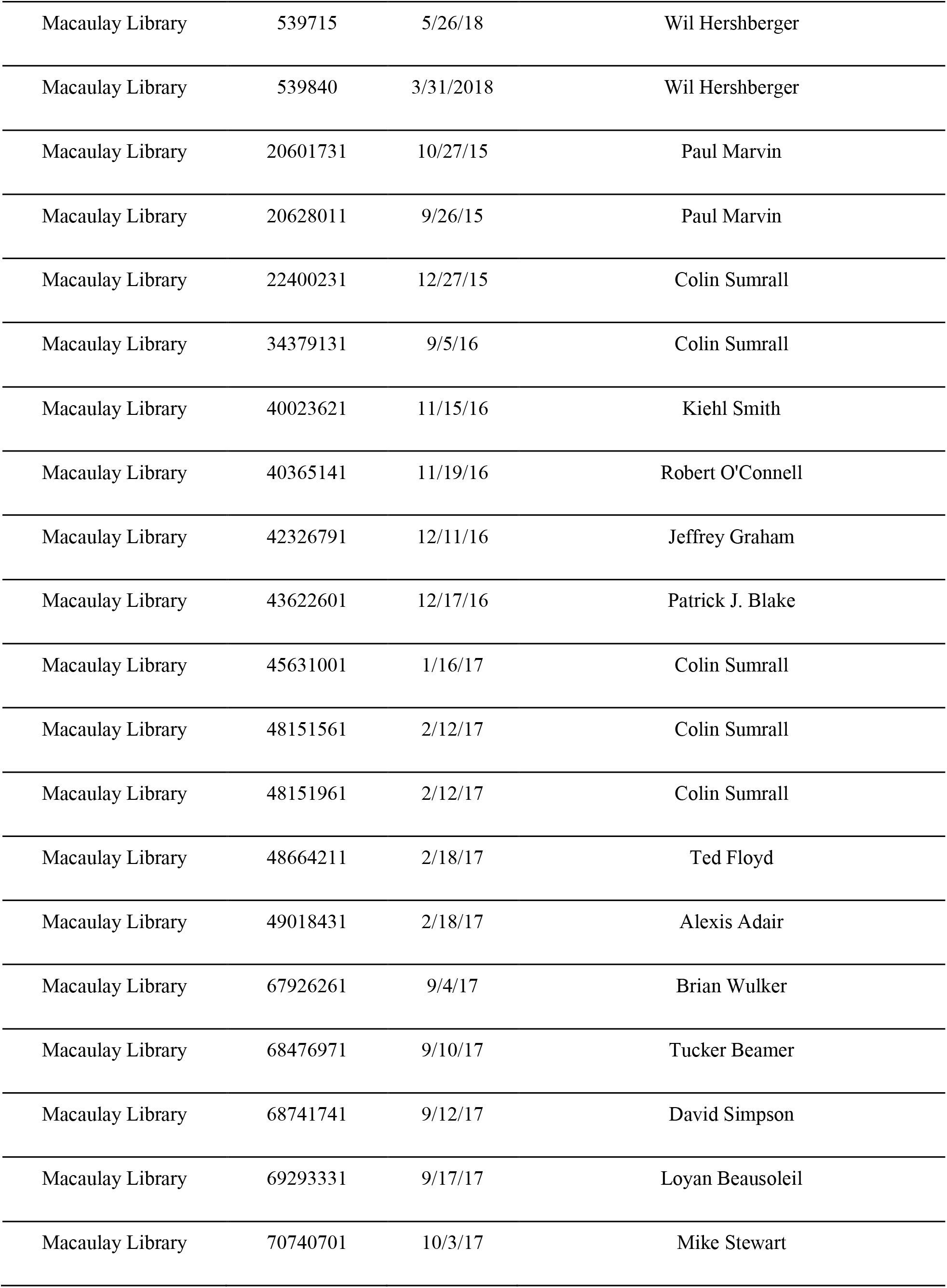
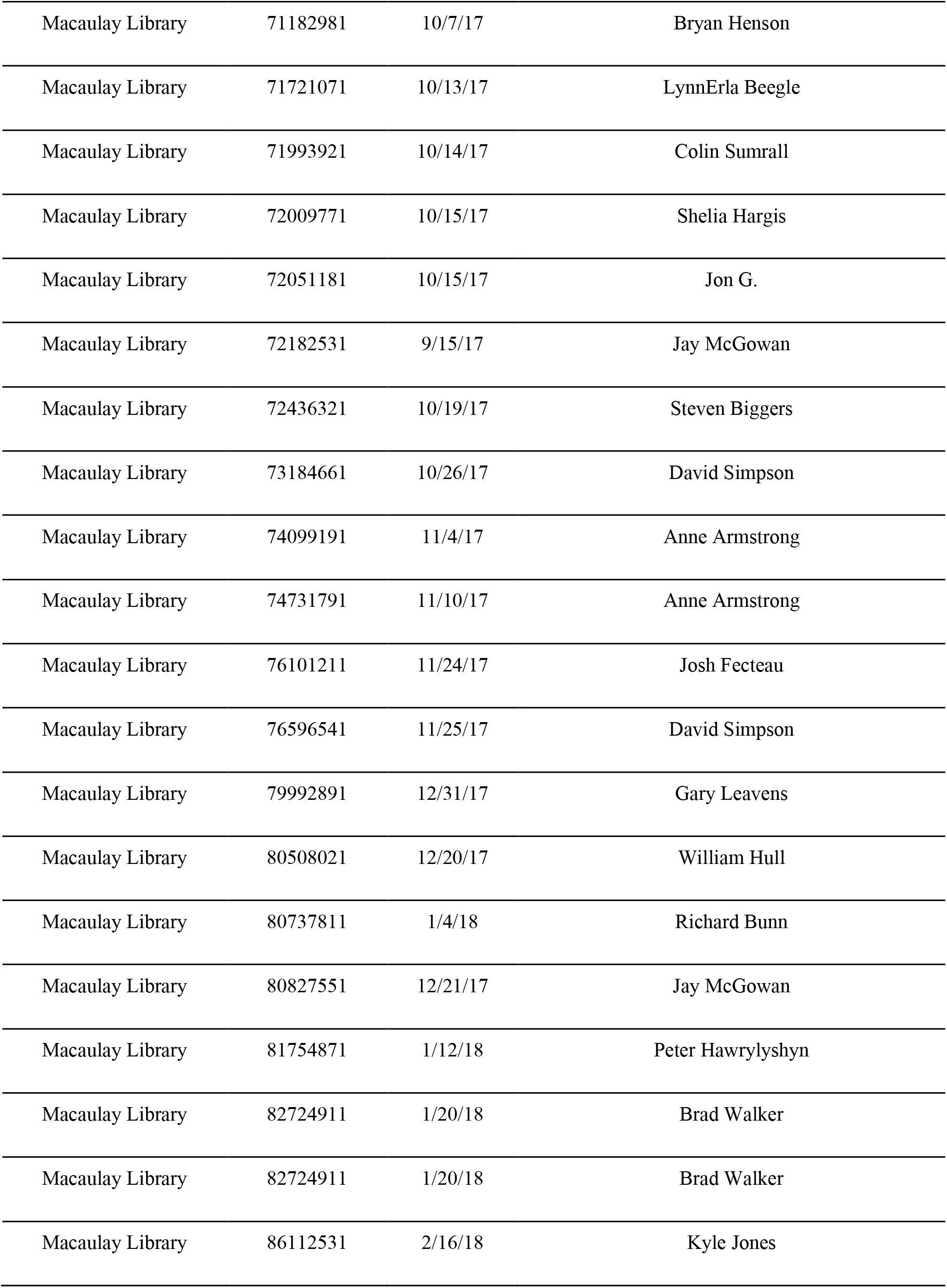
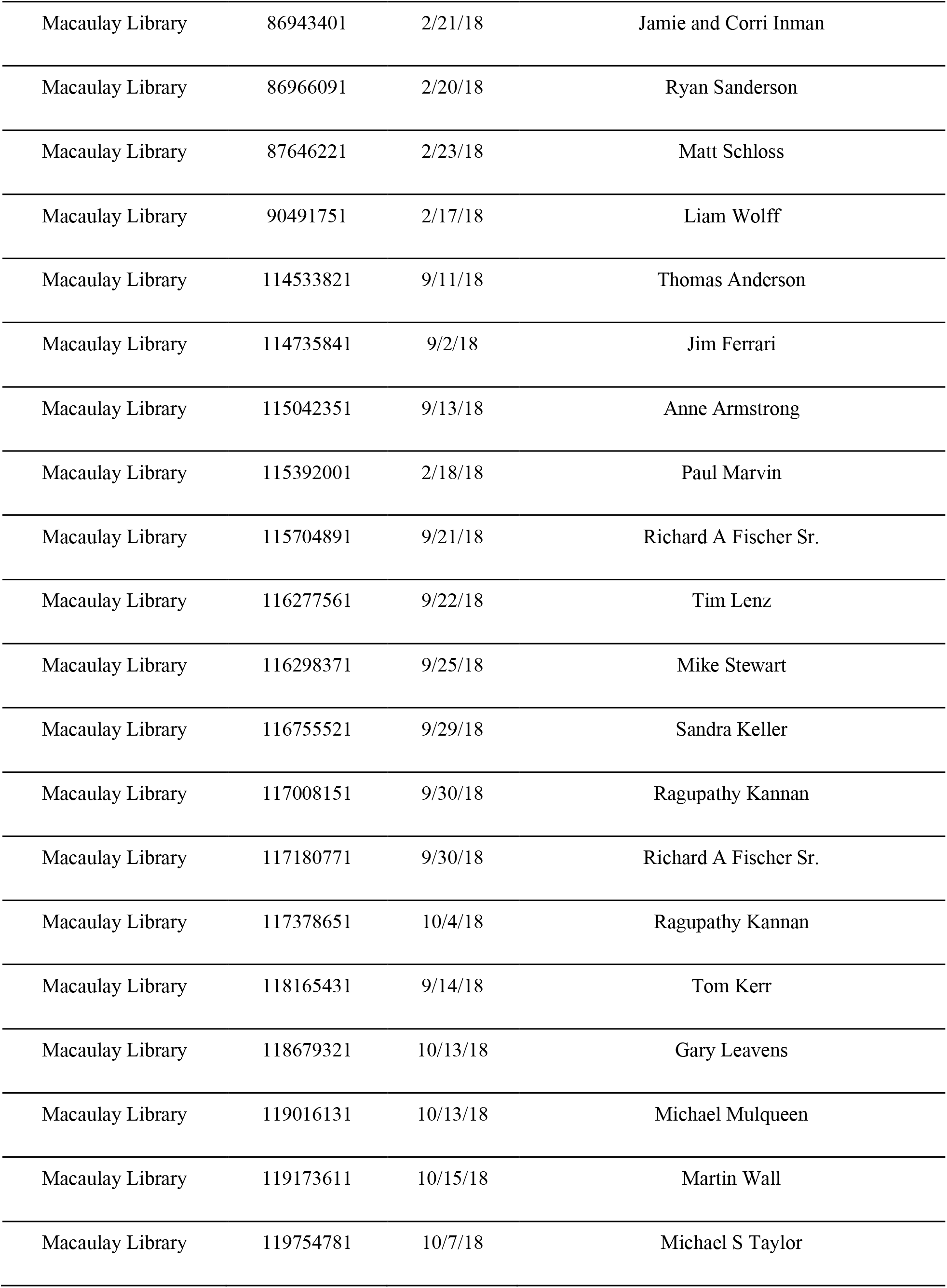
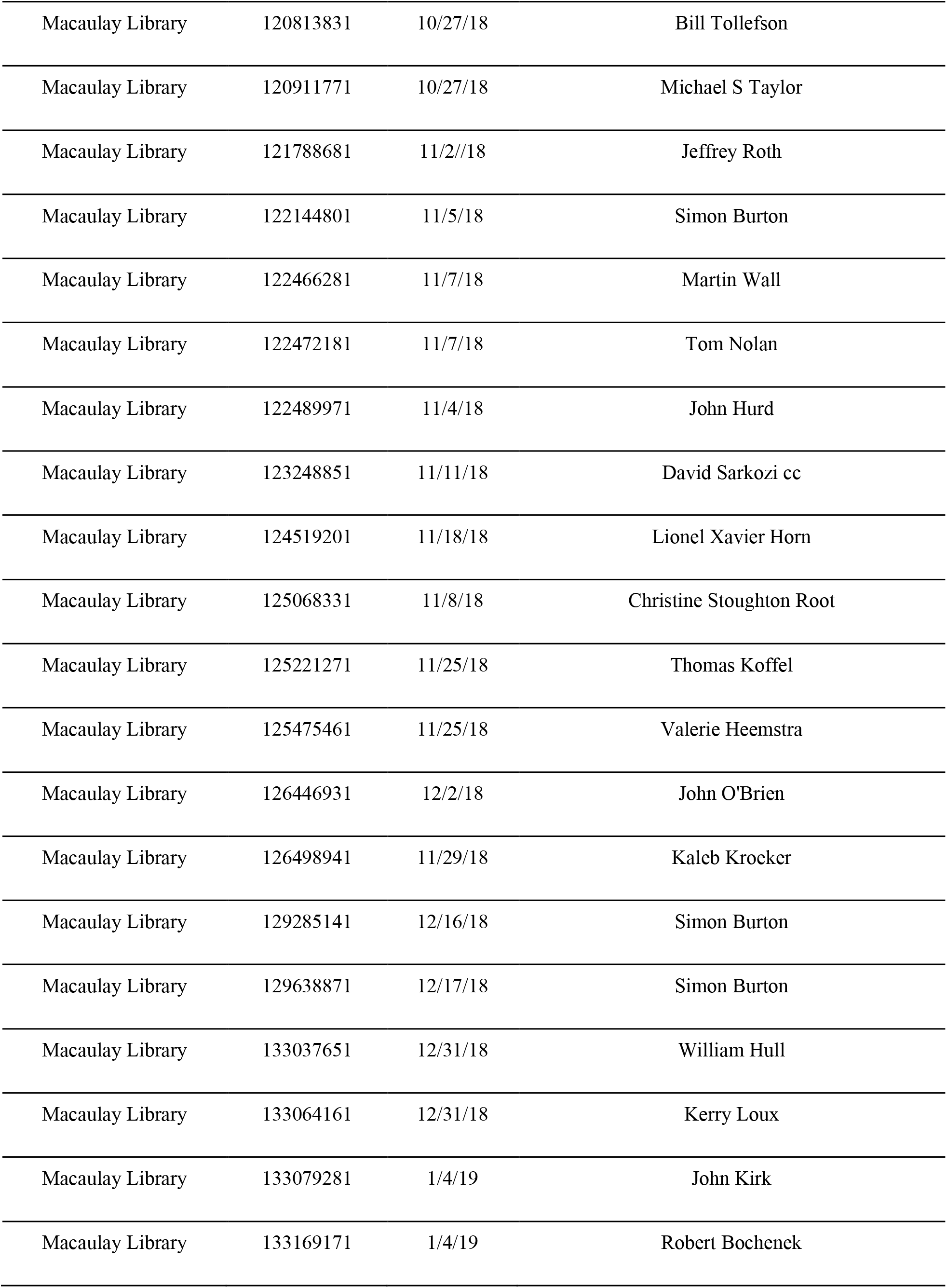
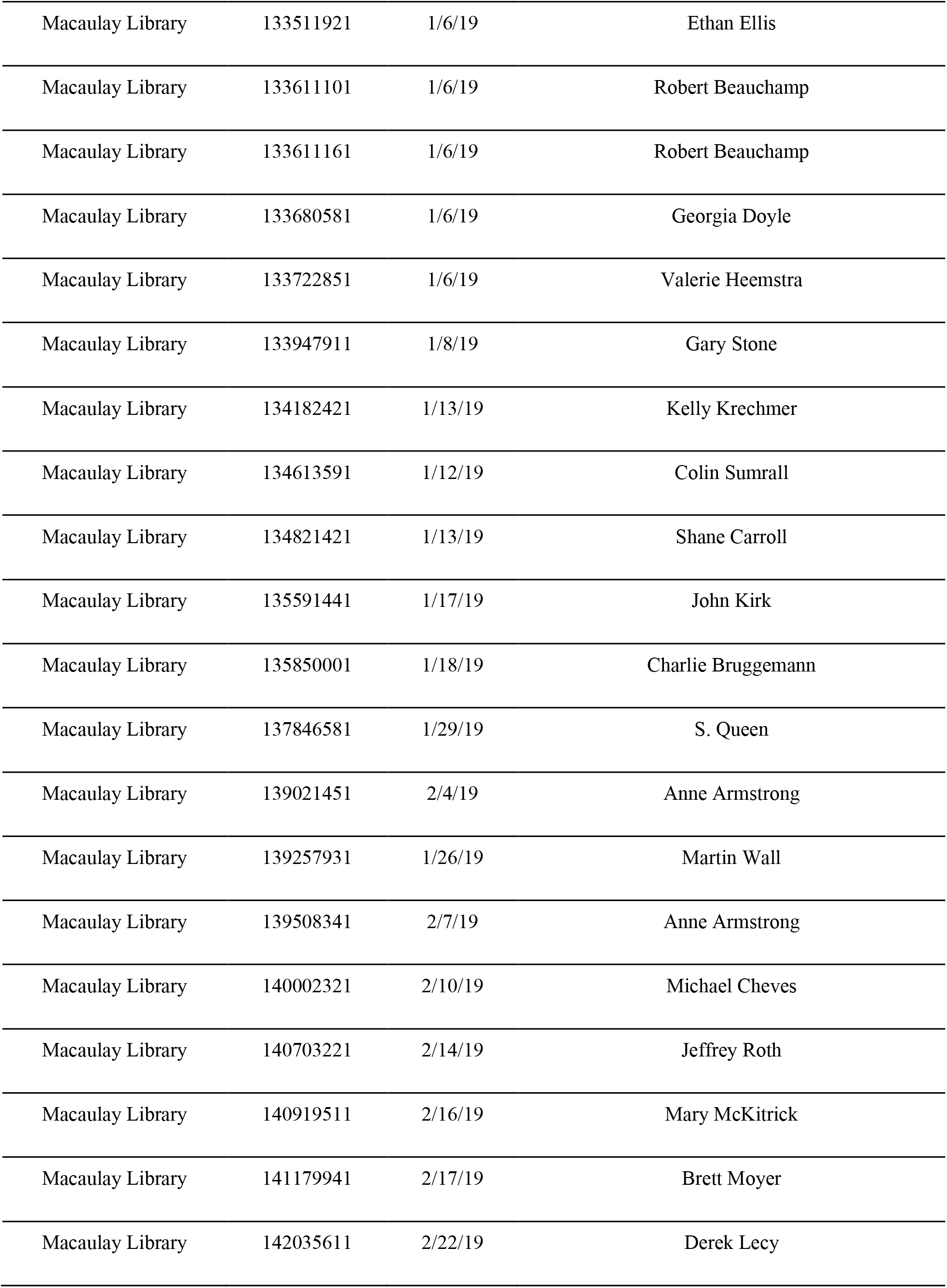
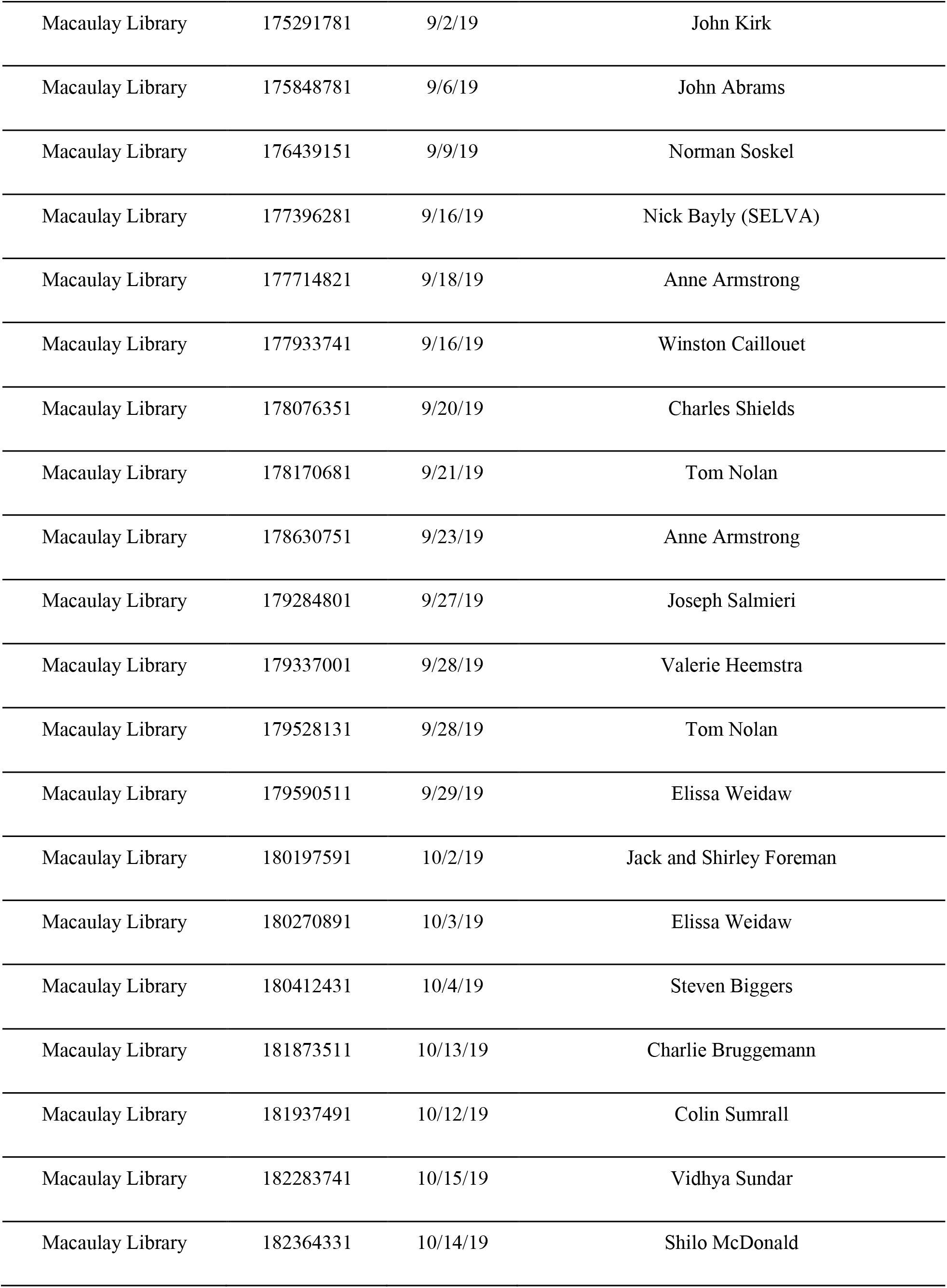
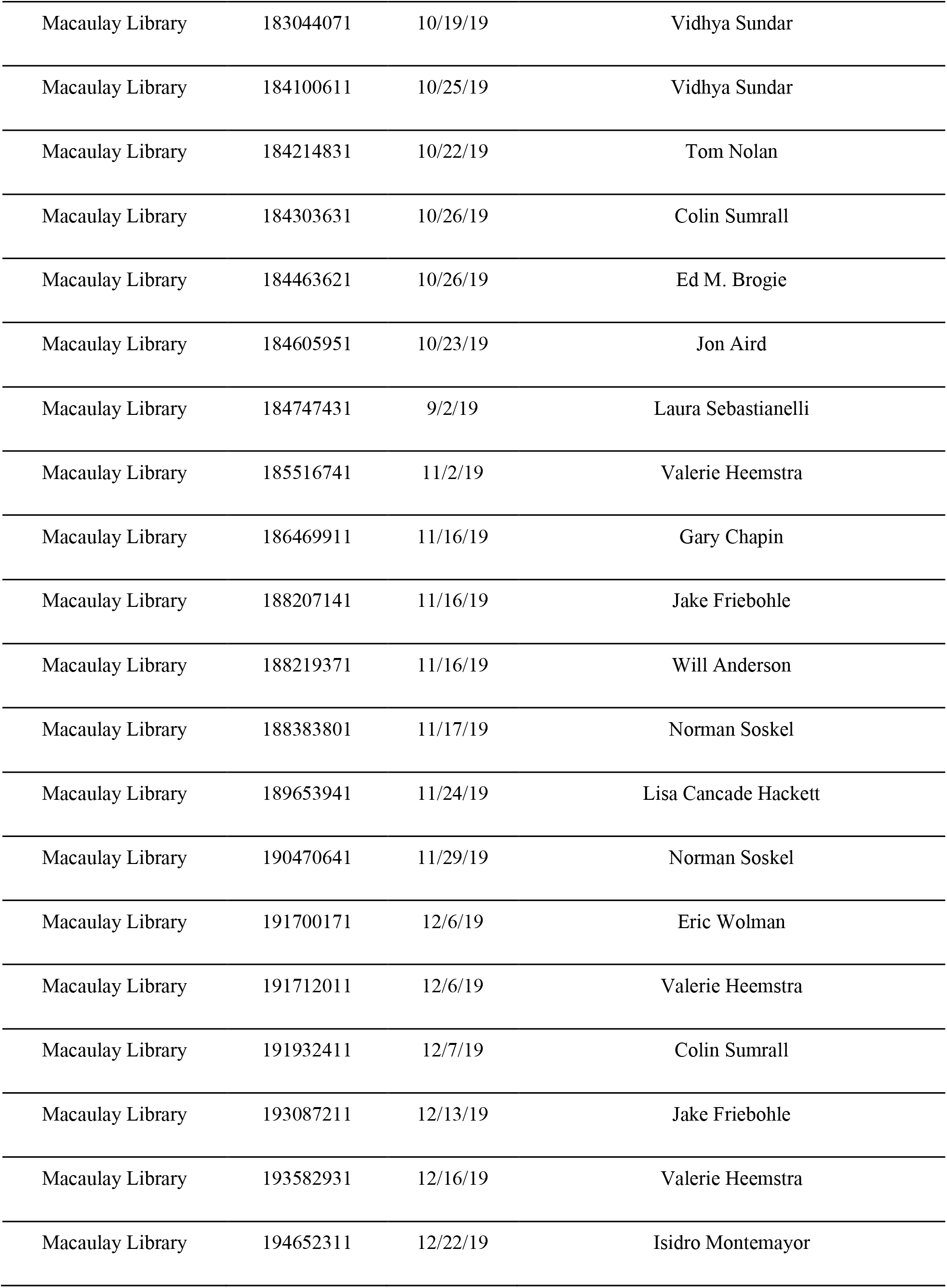
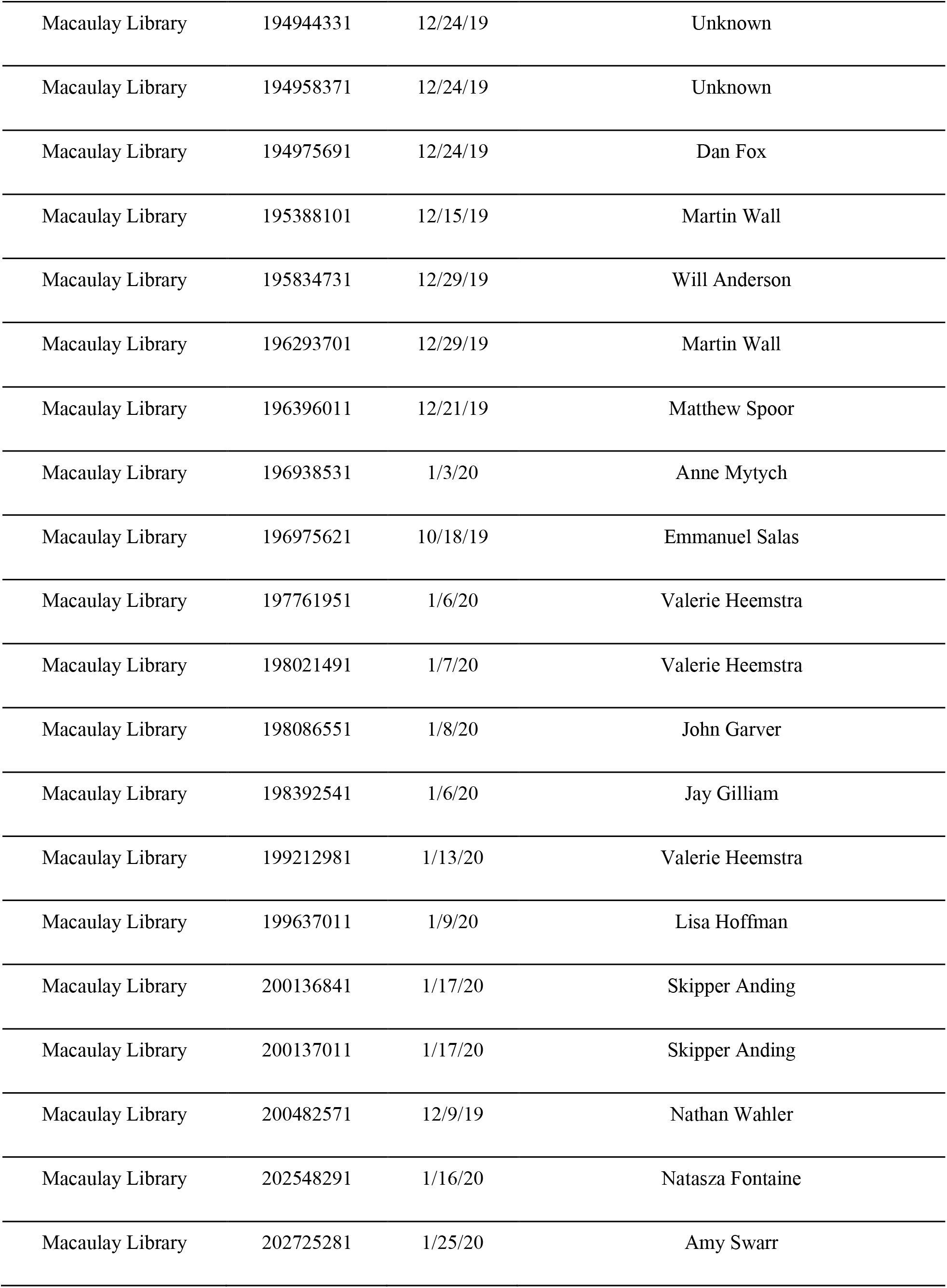
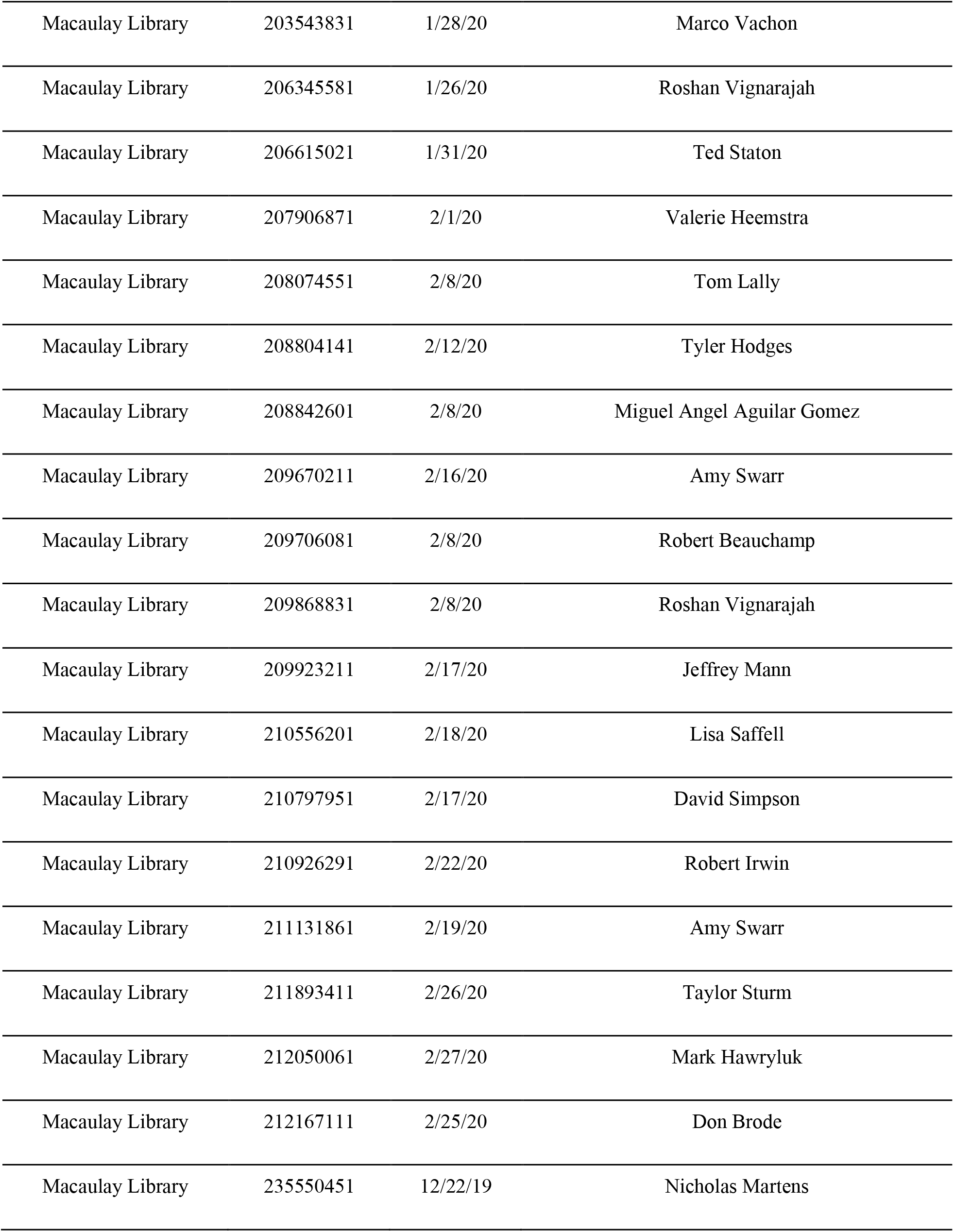
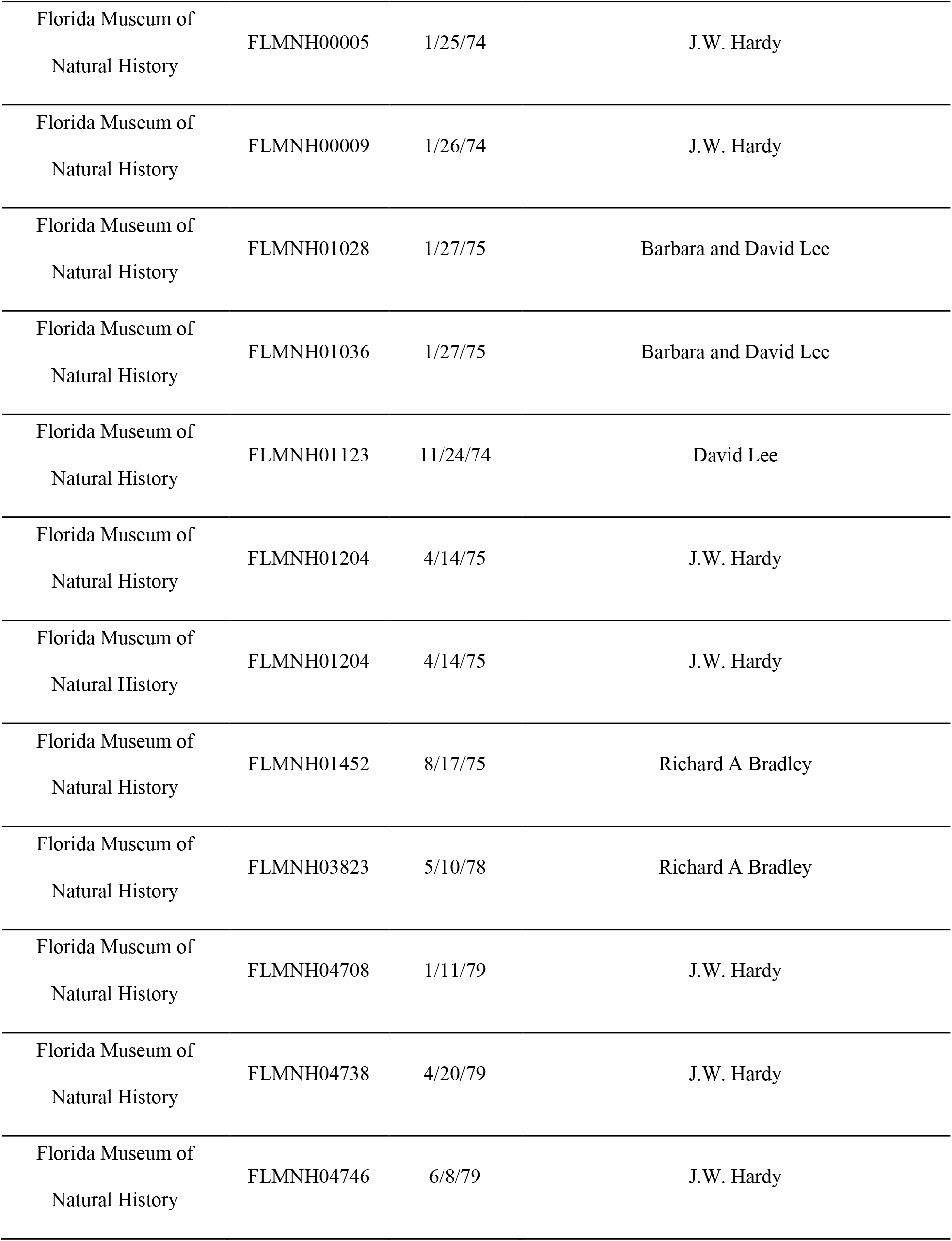
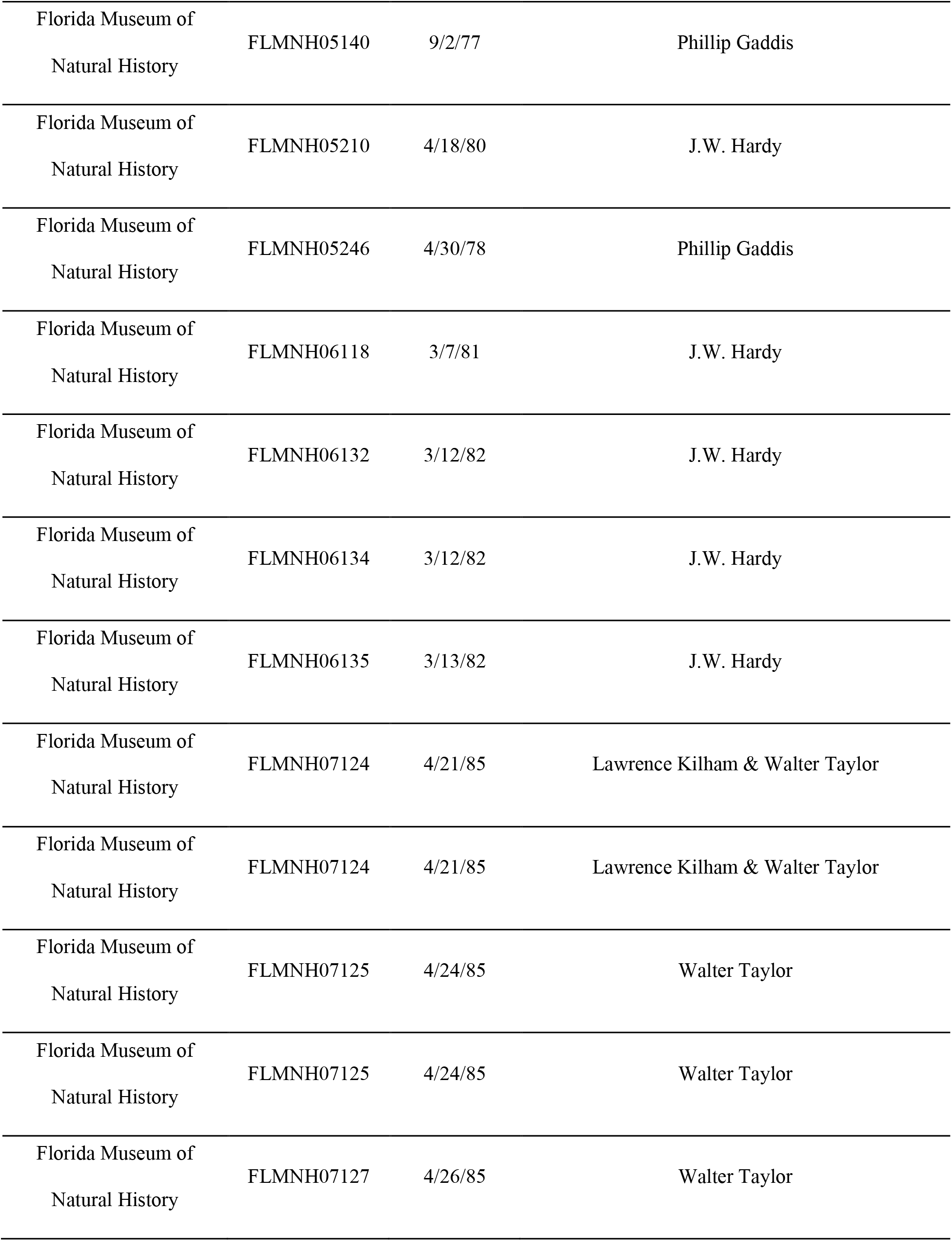
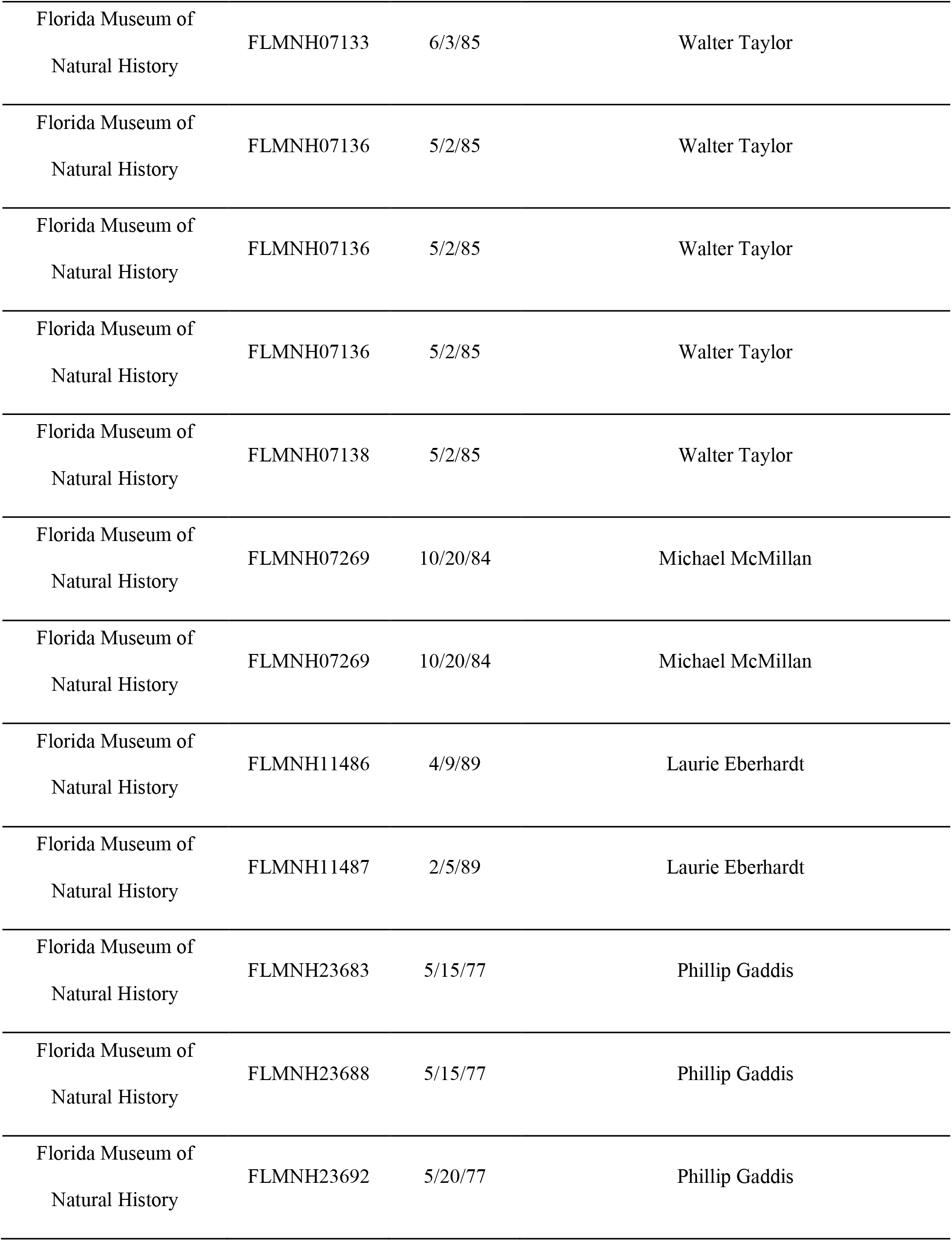
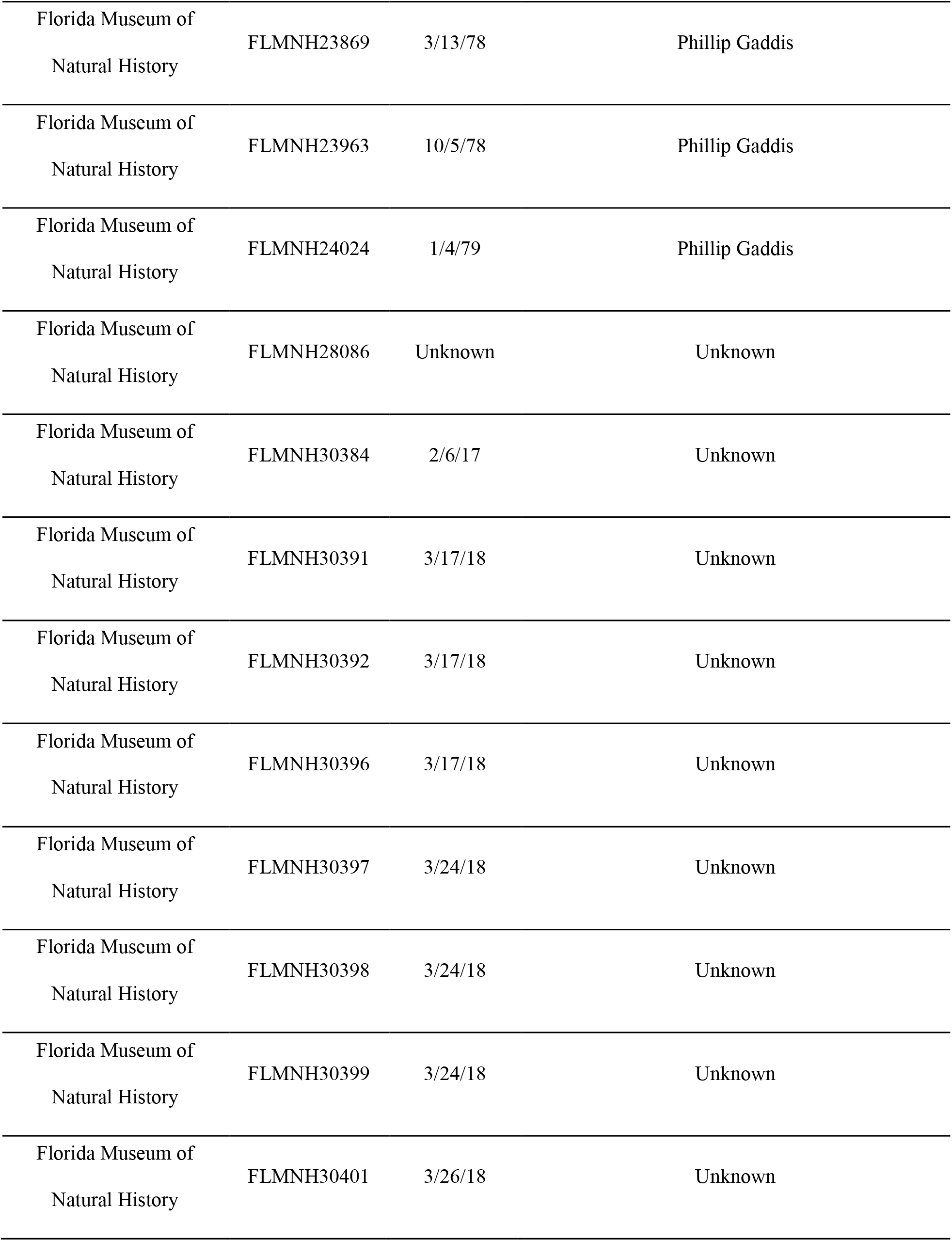
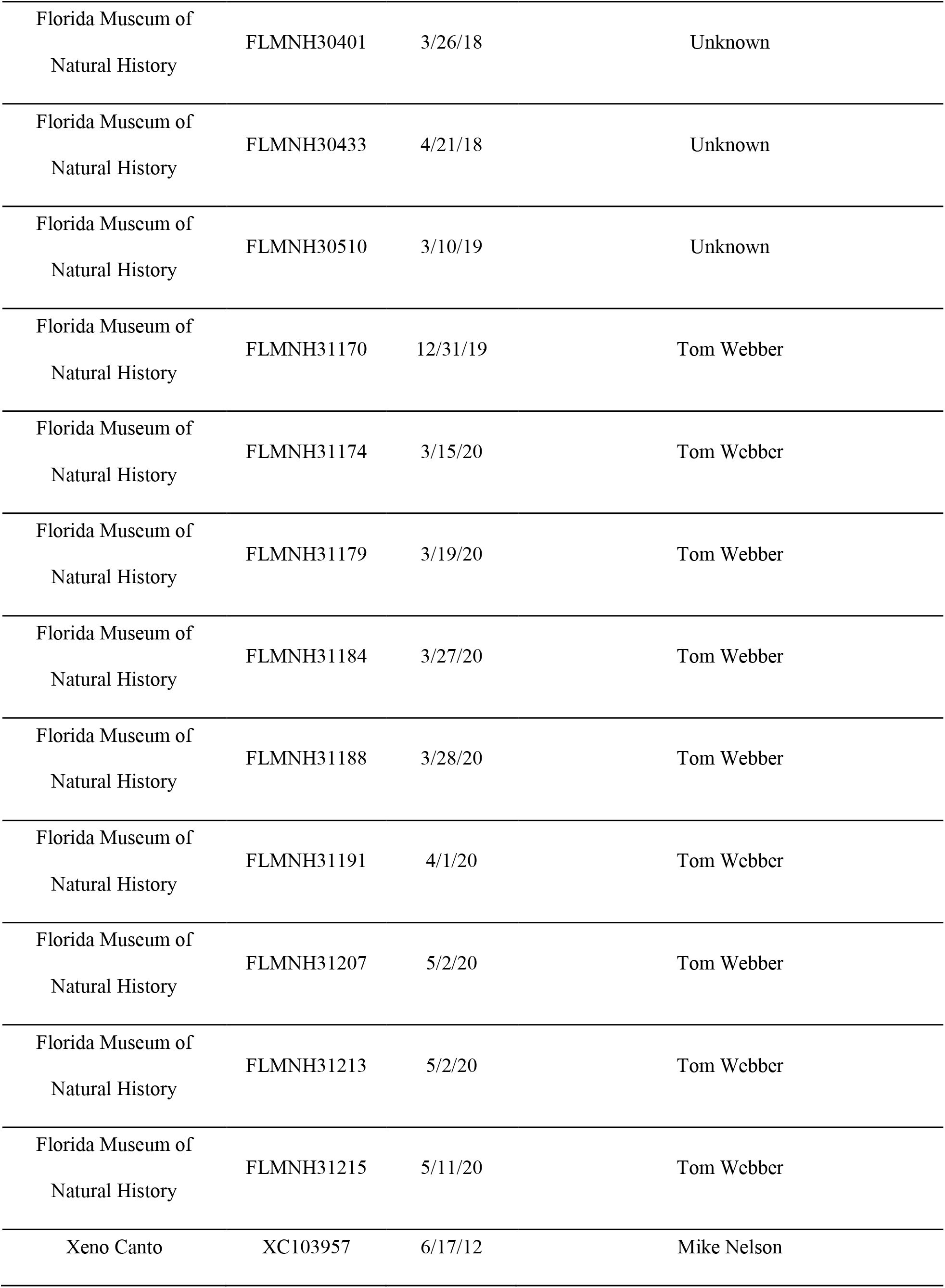
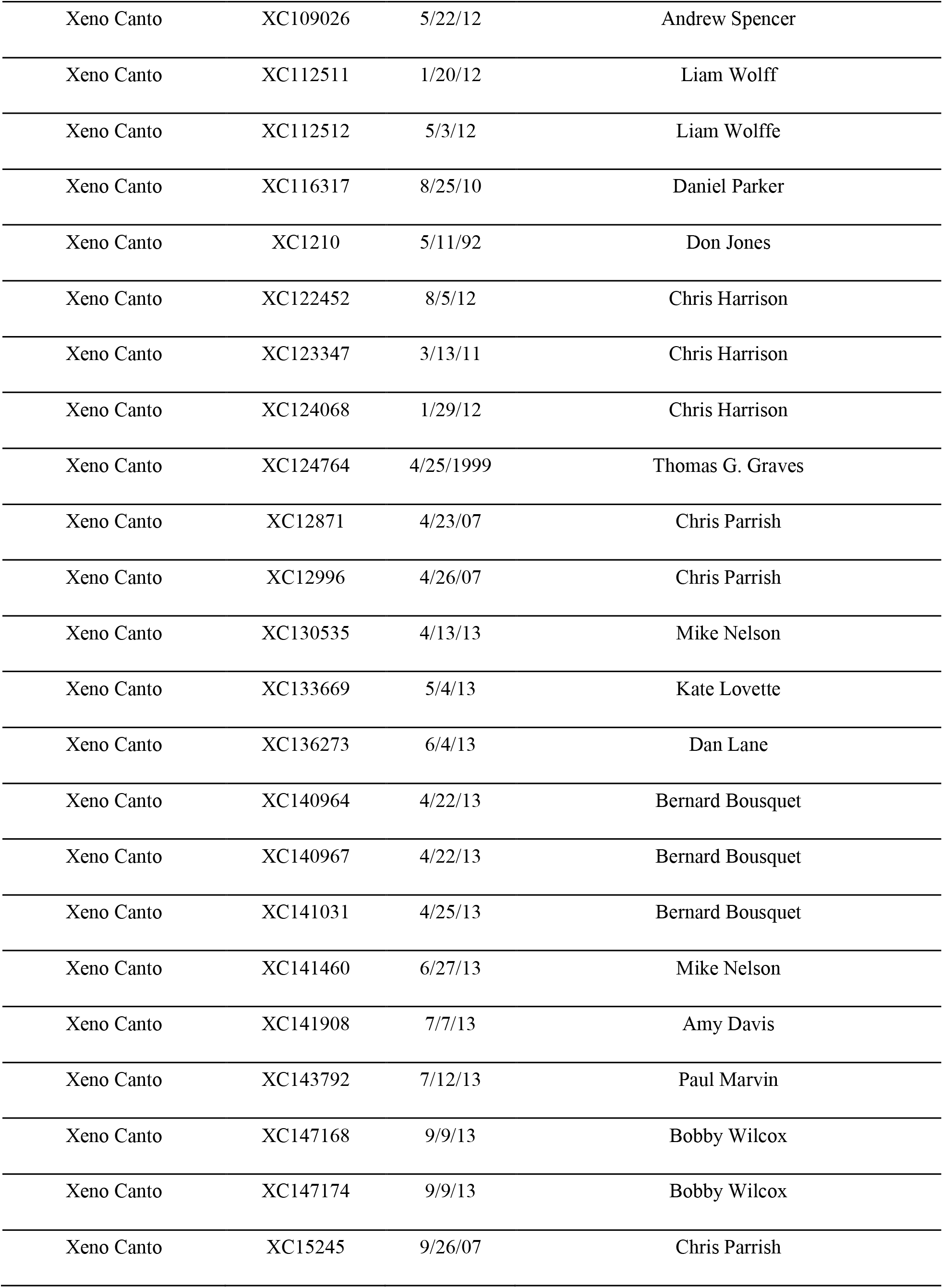
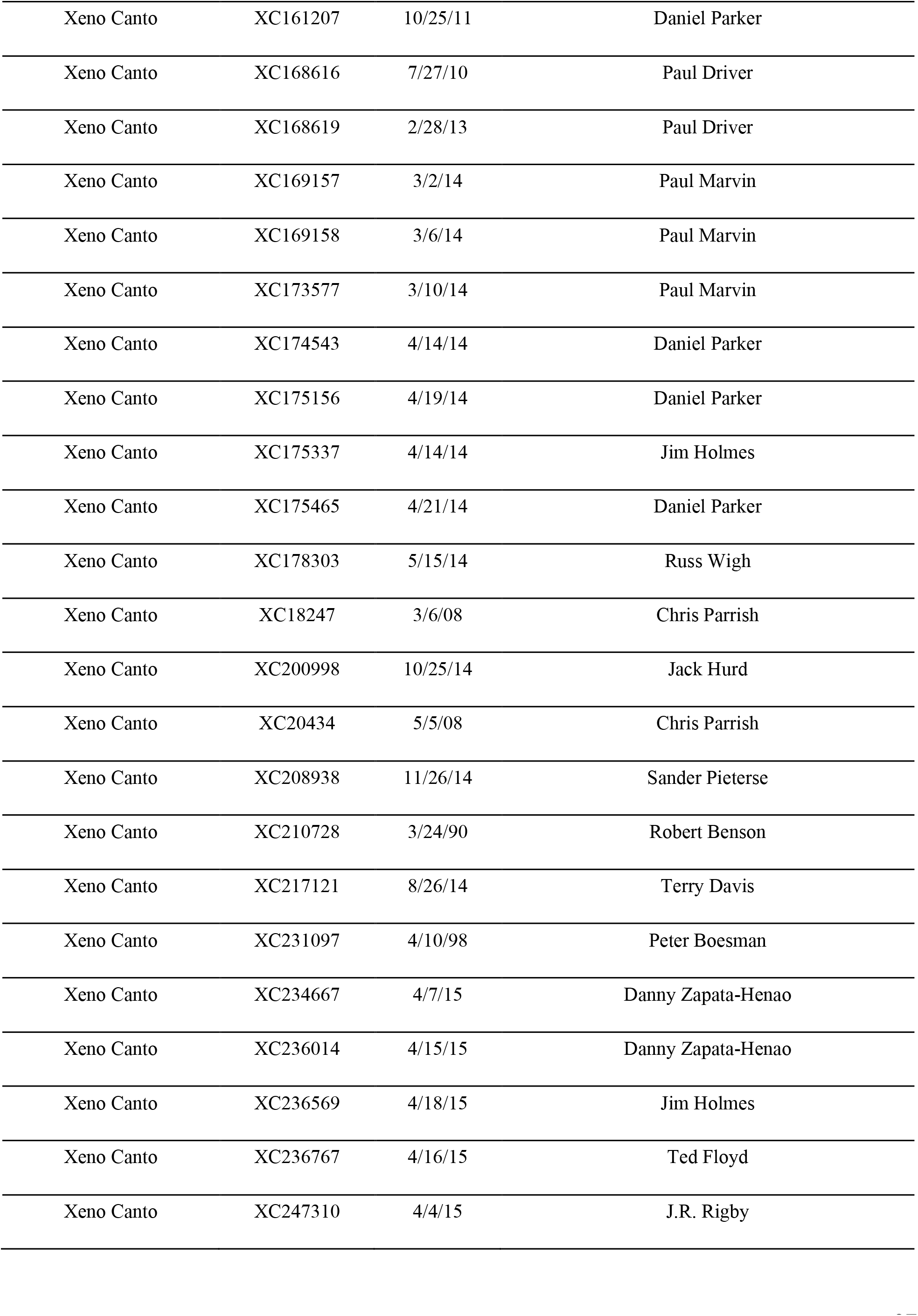
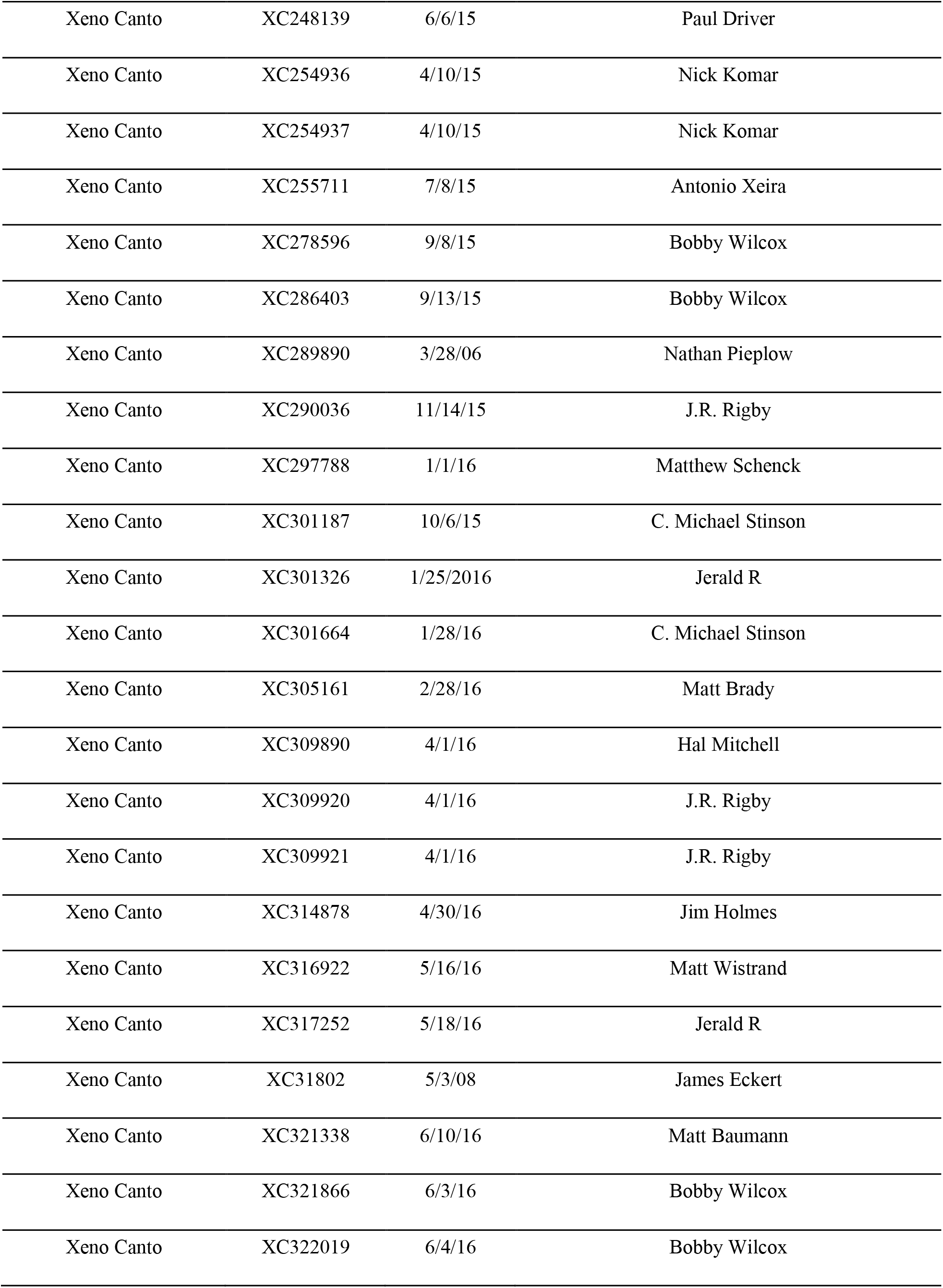
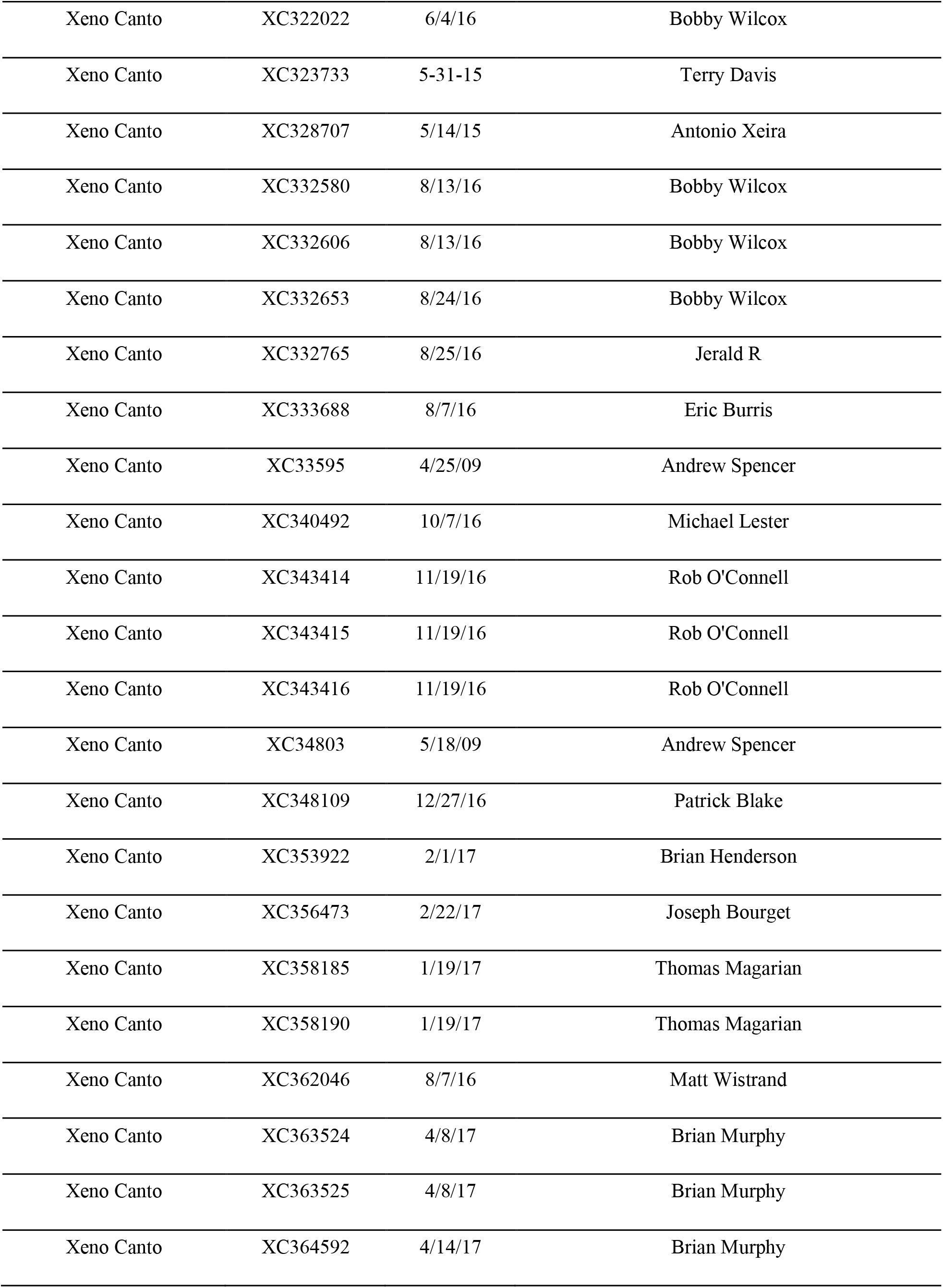
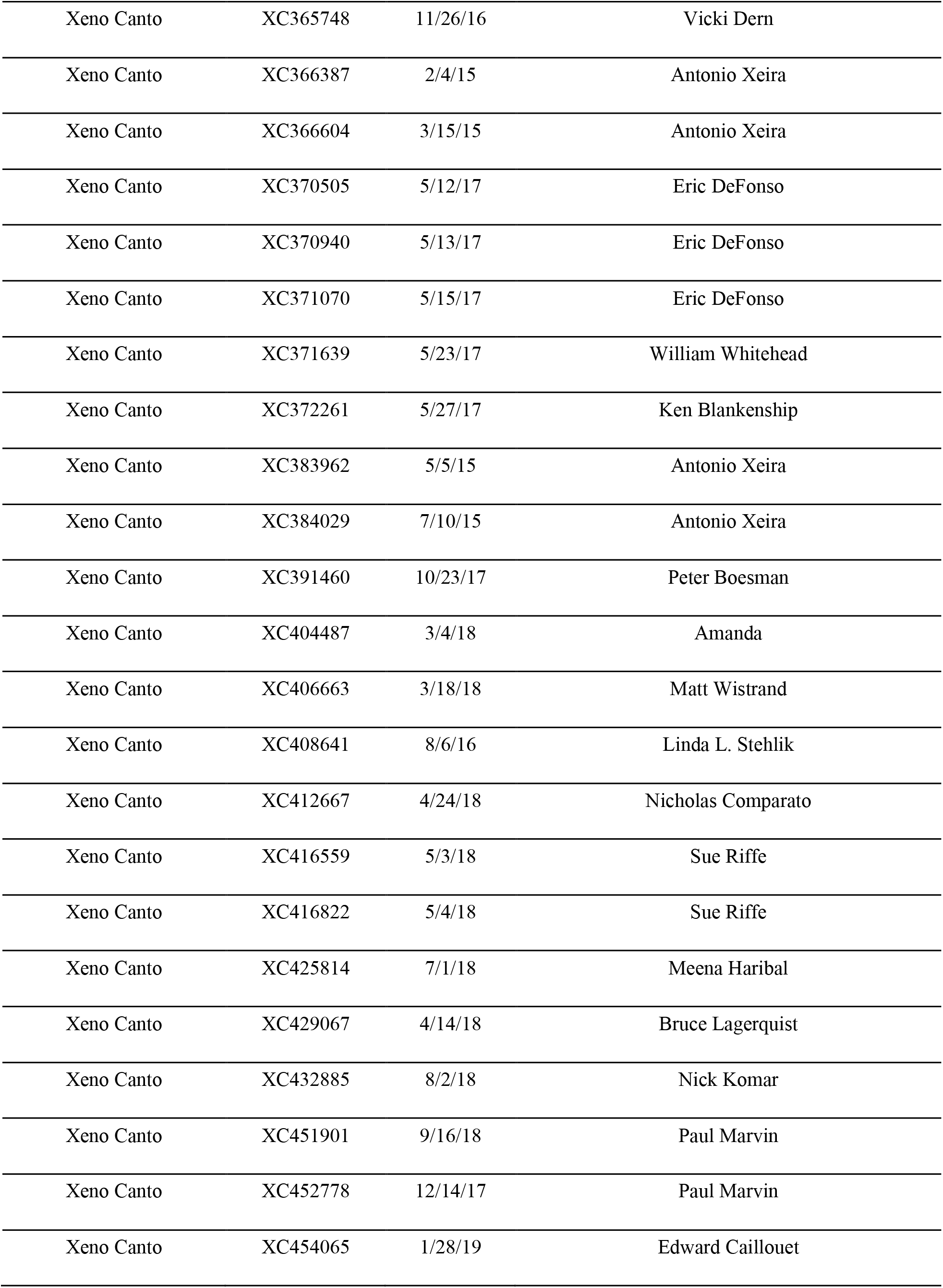
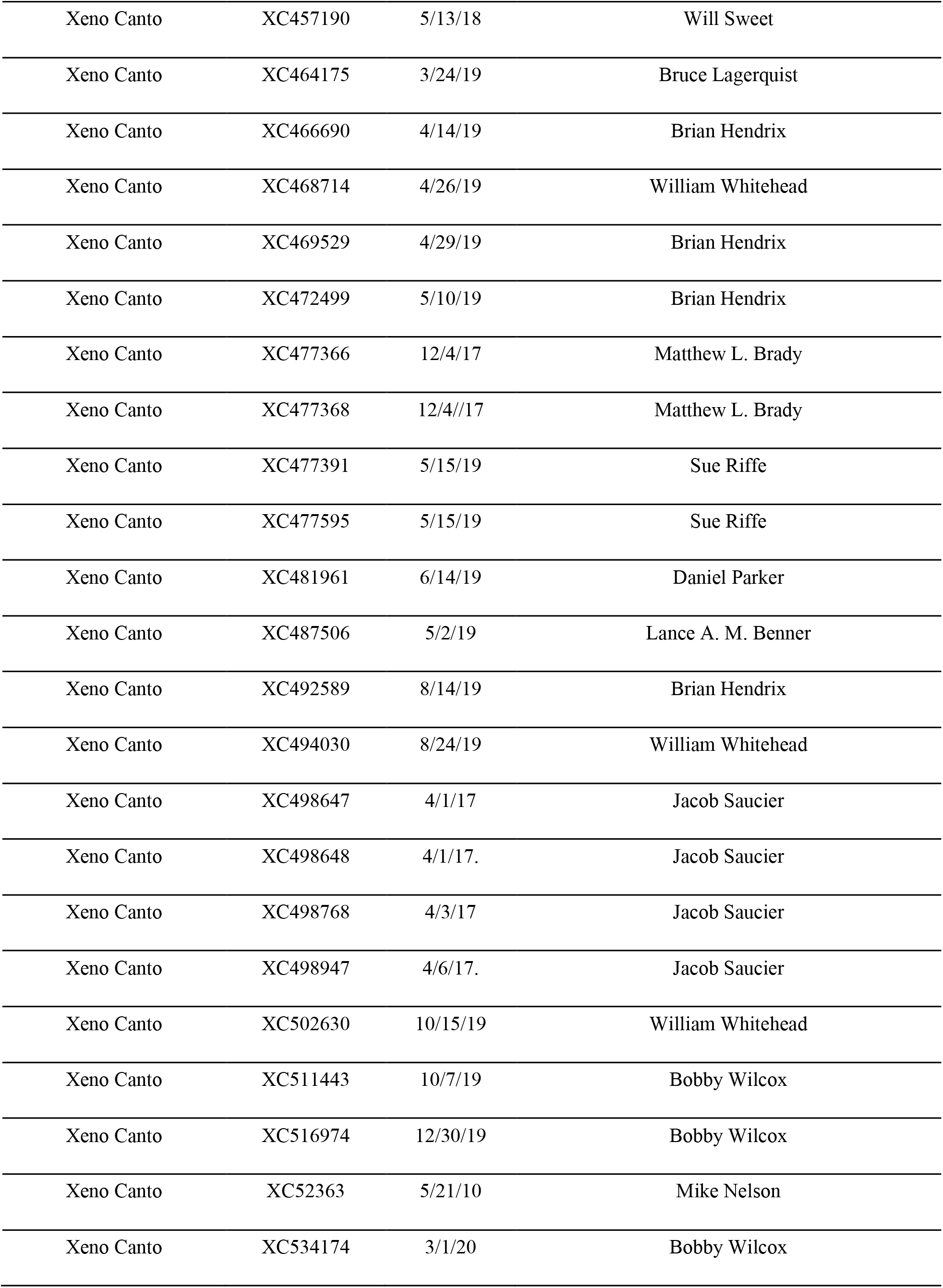
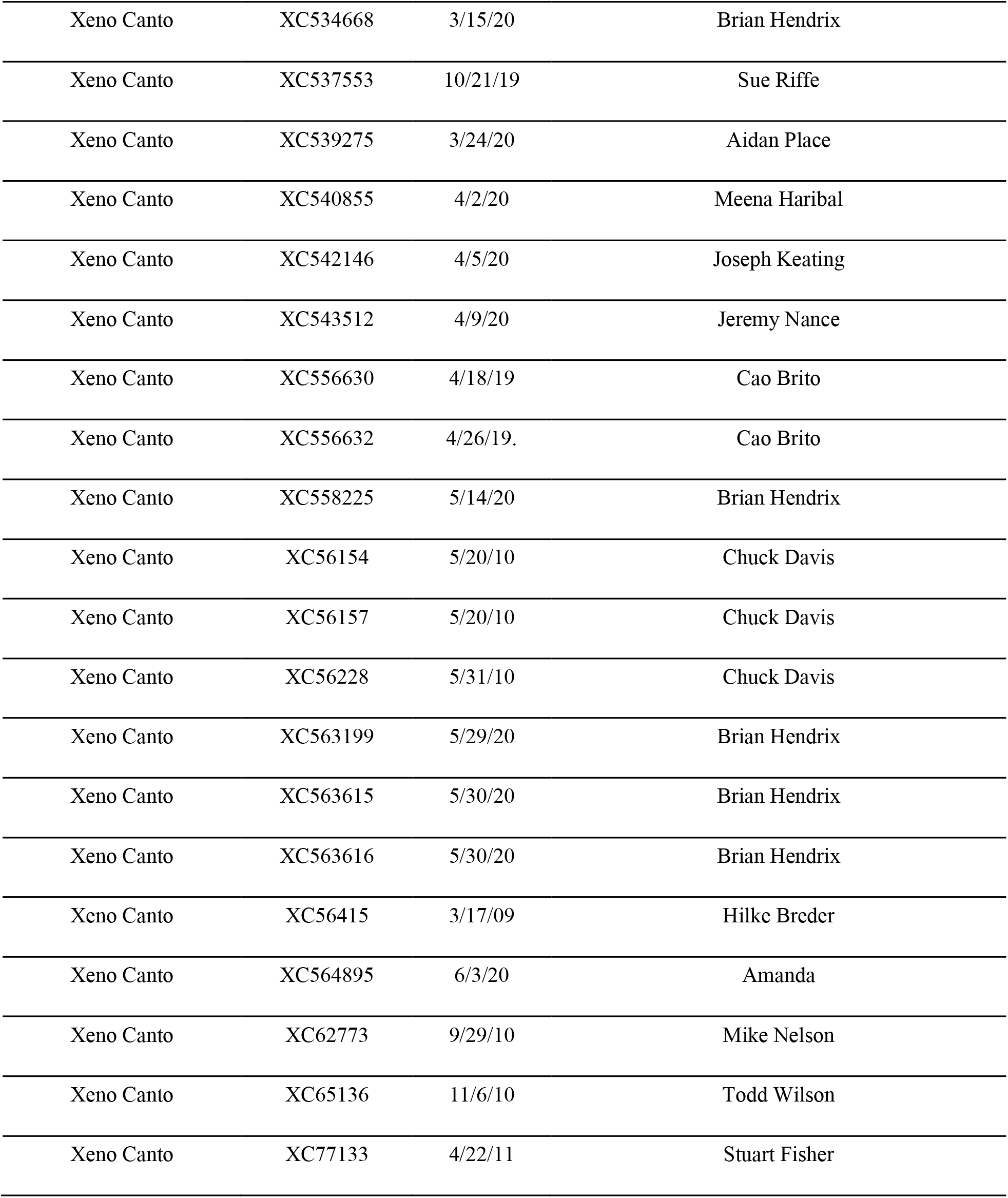
Reference information for the recordings obtained from online databases.

**Supplementary Table 2S:** Results of linear mixed model analysis for song measurements comparing breeding (N = 264) and nonbreeding (N = 177) populations when transitional months between seasons were removed (February and September). All means are presented ± 1 SEM. Bolded values identify P < 0.05.

| Song Measurement | Breeding Mean | Nonbreeding Mean | $X^2_I$ | *P-value* |
| --- | --- | --- | --- | --- |
| **Song Length (sec)** | **1.69±0.02** | **1.57±0.02** | **10.35128** | **<0.05** |
| Maximum Frequency (Hz) | 5464.60±61.63 | 5475.05±75.98 | 0.04127943 | 0.84 |
| Minimum Frequency (Hz) | 1812.73±16.85 | 1801.41±21.16 | 3.670049 | 0.06 |
| Frequency Bandwidth (Hz) | 3651.86±62.64 | 3673.64±78.19 | 0.4214665 | 0.52 |
| Number of Inflection Points | 96.95±1.74 | 95.54±2.29 | 0.1506212 | 0.7 |
| Maximum Entropy | 4.85±0.02 | 4.91±0.03 | 0.3621236 | 0.55 |
| **Number of Syllables/Song** | **4.75±0.09** | **4.28±0.09** | **8.285182** | **<0.05** |
| Number of Notes/Syllable | 4.90±0.07 | 4.86±0.08 | 0.007514044 | 0.93 |
| Trill Rate (syllables/sec) | 3.04±0.06 | 3.16±0.06 | 5.571908 | 0.81 |

